# Phytopathogenic cyclic glucohexadecaose from an inverting transglycosylase

**DOI:** 10.1101/2023.11.26.568766

**Authors:** Sei Motouchi, Shiro Komba, Hiroyuki Nakai, Masahiro Nakajima

**Affiliations:** Department of Applied Biological Science, Faculty of Science and Technology, Tokyo University of Science, 2641 Yamazaki, Noda Chiba 278-8510, Japan; Division of Food Processing and Biomaterials Biomaterials Development Group, Institute of Food Research, National Agriculture and Food Research Organization, 2-1-12, Kannondai, Tsukuba, Ibaraki 305-8642, Japan; Faculty of Agriculture, Niigata University, 8050 Ikarashi 2-no-cho, Nishi-ku, Niigata 950-2181, Japan

## Abstract

*Xanthomonas* species contain numerous notoriously well-known plant pathogens. Among various pathogenic factors, the role of α-1,6-cyclized β-1,2-glucohexadecaose (CβG16α) produced by *Xanthomonas campestris* pv. *campestris* was shown previously to be vital for infecting model organisms *Arabidopsis thaliana* and *Nicotiana benthamiana*. However, enzymes responsible for biosynthesising CβG16α are essentially unknown, which limits the generation of agrichemicals that inhibit CβG16α synthesis. In this study, we discovered that OpgD from *X. campestris* pv. *campestris* converts linear β-1,2-glucan to CβG16α. Structural and functional analyses revealed that OpgD from *X. campestris* pv. *campestris* possesses an anomer-inverting transglycosylation mechanism, which is unprecedented among carbohydrate-active enzymes. The discovery of this unprecedented glucan-generating mechanism reveals a new foundation for the enzymatic synthesis of carbohydrates. Furthermore, identifying CβG16α synthase highly conserved in *Xanthomonas* provides a broadly adaptable drug target for new-genre agrichemicals that overcome antimicrobial-resistant bacterial issues.

---

*Xanthomonas* species are world-notorious plant pathogens that cause diseases in more than 400 different plant hosts such as rice, wheat, tomato, pepper, cabbage, cassava, banana and bean^1^. Although antimicrobial agrichemicals have been used mainly for avoiding virulence, antimicrobial-resistant bacteria that emerge by natural selection are a severe problem in agriculture. New-genre agrichemicals that inhibit pathogenicity without being invalidated by natural selection are in demand.

α-1,6-Cyclized β-1,2-glucohexadecaose (CβG16α) produced by *Xanthomonas campestris* pv*. campestris*^2^ is a potential broad-adaptable inhibition target for new-genre agrichemicals. CβG16α is vital for the pathogenicity toward model plants *Arabidopsis thaliana* and *Nicotiana benthamiana*, despite the non-production of CβG16α not affecting the fertility of *X. campestris* pv*. campestris*^3^. In detail, CβG16α suppresses the expression of PR (pathogenesis-related) proteins and callose accumulation in plants^3^. However, key enzymes responsible for the biosynthesis of CβG16α remains unknown.

CβG16α is classified into osmo-regulated periplasmic glucans (OPG)^4^. There are major patterns in OPGs with β-1,2-linked glucosyl backbone, namely α-1,6-cyclized β-1,2-glucooligosaccharide, cyclic β-1,2-glucan (CβG) and β-1,2-glucooligosaccharide with β-1,6-glucose side chains (LβG-6β)^4^. Phenotypes of various mutants with genes related to the biosynthesis of OPG mutated (not limited to “OPG synthesis” itself) have been analysed. Most of these mutants showed significantly different phenotypes from the wild-type species, such as lack of pathogenicity (*X. campestris* pv. *campestris*, *Brucella abortus*, *Agrobacterium tumefaciences*, *Pseudomonas syringae*, *Dickeya dadnantii*, *Salmonella enterica* sv. *typhimurium*) or symbiotic ability (some *Rhizobiaceae*)^4–11^. Although the enzyme synthesizing CβG has been identified, many other enzymes responsible for biosynthesizing carbohydrate backbones of OPG have not been explored^4^. For example, *Escherichia coli* synthesizes a LβG-6β-type OPG. However, even in *Escherichia coli*, a well-known model organism, the biochemical functions of enzymes required after elongation of linear β-1,2-glucan were unknown.

Recently, Motouchi et al. identified a group of OPG-related proteins with unknown biochemical functions (MdoG superfamily) as a phylogenetically new glycoside hydrolase (GH) family, GH186. OpgD from *E. coli* (EcOpgD), a GH186-establishing enzyme, was elucidated to be a β-1,2-glucanase (SGL) (Supplementary Note 1) that regulates the chain length of LβG-6β^12^. A remarkable feature in GH186 is the amino acid sequence diversity of a region vital for the unique hydrolytic reaction mechanism of EcOpgD. This observation indicates diverse functions and reaction mechanisms among GH186 enzymes which are distributed mainly among α, β and γ-proteobacteria, including *Xanthomonas campestris* pv. *campestris*^12^.

In this report, we discovered that XccOpgD, a GH186 homologue from *X. campestris* pv. *campestris* alone converts linear β-1,2-glucan into CβG16α specifically. Structural analysis of XccOpgD revealed an unprecedented reaction mechanism, anomer-inverting transglycosylation, revolutionising the long-believed patterns of the reaction mechanisms among GH enzymes. The results further indicated diverse OPG biosynthesis-related functions exist other than CβG16α and LβG-6β synthesis by enzymes of GH186, leading to the concept of a broadly adaptable agrichemical free of antimicrobial resistance.

## Results

### Identification of the reaction product by XccOpgD

Recombinant XccOpgD fused with a His_6_-tag at the C-terminus was produced using *E. coli* as a host and purified successfully. XccOpgD exhibited activity toward linear β-1,2-glucans to produce glucans with degrees of polymerization (DPs) adjusted, and no by-product was detected by TLC (Fig. 1a). The product was not hydrolysed by treatment with a β-glucosidase from *Bacteroides thetaiotaomicron* (BtBGL)^13^, an exo-type enzyme (Fig. 1a), indicating that the product is cyclized or modified at the non-reducing end. Electrospray ionization-mass spectrometry (ESI-MS) analysis revealed that the DP of the product was 16, and the molecular mass was lower than linear β-1,2-glucan by 18 mass units, indicating the product is a cyclic form (Fig. 1b, Extended Data Fig. 1 and Extended Data note 2). One- and two-dimensional NMR analysis was performed to determine the chemical structure of the product. Only one α-1,6-glucosidic bond was identified, with all other bonds being the β-1,2-glucoside type (Fig. 1c; see Extended Data Table 1, 2, and Extended Data Fig. 2 and Supplementary Data 1 for further data). Thus, the product was identified as α-1,6-cyclized β-1,2-glucan consisting of 16 glucose units (α-1,6-cyclized β-1,2-glucohexadecaose, CβG16α) without side chains.

**Fig. 1.**
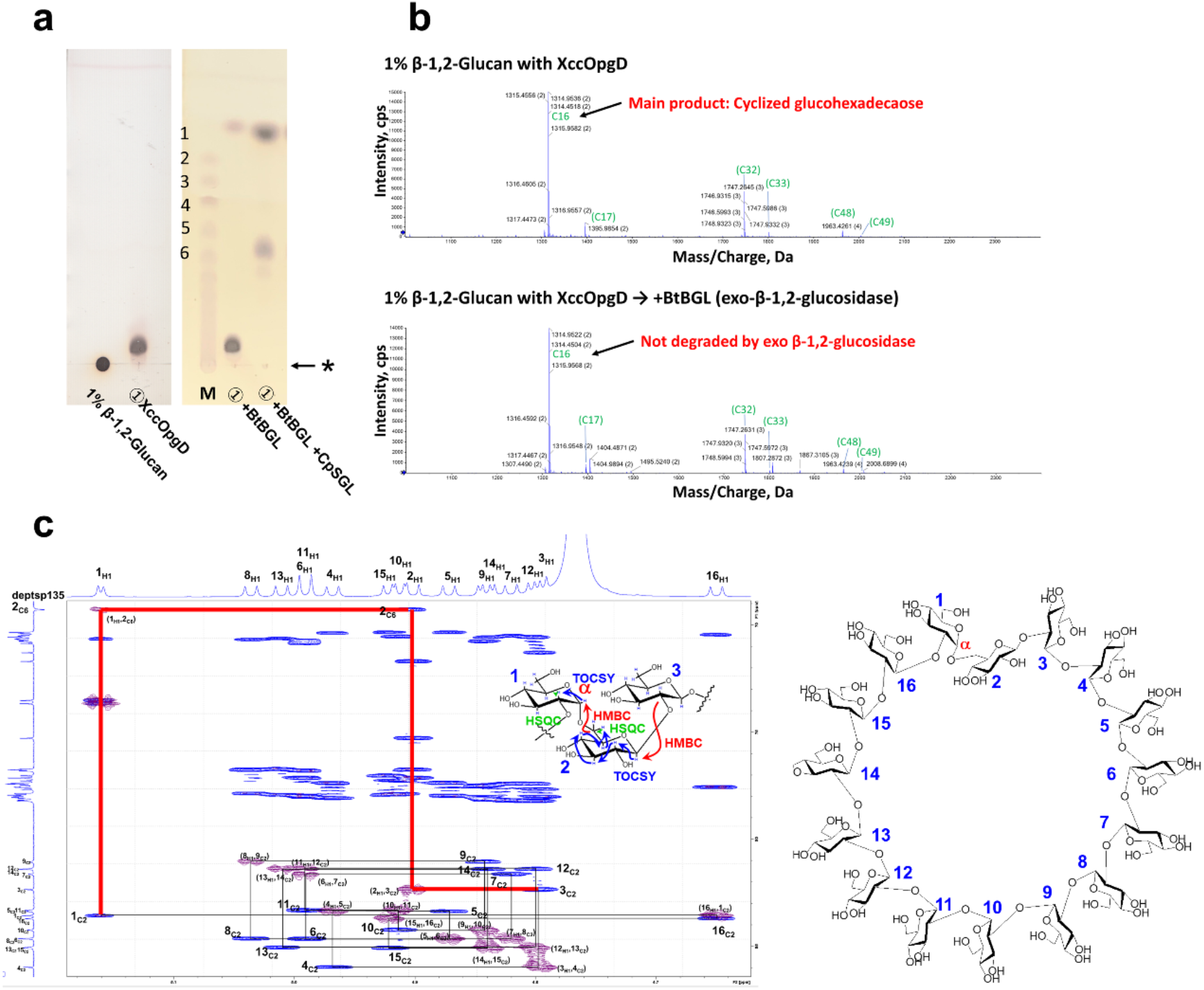
Identification of the main oligosaccharide produced by XccOpgD. **a**, TLC analysis of the product produced by XccOpgD. Lane M, a linear β-1,2-glucooligosaccharide marker prepared using 1,2-β-oligoglucan phosphorylase from *L. innocua*. DPs of linear β-1,2-glucooligosaccharides are shown on the left side of the TLC plates. The origins of the TLC plates are shown as horizontal arrows denoted by asterisks. ①, Linear β-1,2-glucan (1%) was incubated with XccOpgD (0.55 mg/mL) at 37 °C for 24 h. ① + BtBGL, BtBGL (the final concentration was 0.091 mg/mL) was added to ① and incubated at 37 °C for 24 h. ① + BtBGL + CpSGL, BtBGL and CpSGL (final concentrations were 0.083 and 0.25 mg/mL, respectively) were added to ① and incubated at 37 °C for 24 h. **b**, Electrospray ionization-mass spectrometry analysis of the reaction products. The peaks are assigned as [M + nNH_4_]^n+^ and the arrow indicates the main product. Green letters and numbers represent forms of compounds (cyclic or linear) and DPs of products, respectively. For example, C16 represents cyclized hexadecaose. Green letters in parentheses are the peaks thought to be derived from cyclic glucohexadecaose. (top) Reaction products released from linear β-1,2-glucan by XccOpgD. (bottom) Reaction products after BtBGL treatment. **c**, NMR analysis of the purified main product released by XccOpgD. Blue and purple peaks represent HSQC-TOCSY and HMBC data, respectively. The numbers of Glc moieties are defined by the blue numbers. Black lines trace from 1_C2_ (2-carbon at the Glc moiety 1) to 3_C2_ (2-carbon at the Glc moiety 3) in the non-reducing end direction. Red lines trace from 3_C2_ to 1_C2_ in the non-reducing end direction.

**Table 1.** Kinetic parameters of XccOpgD for linear β-1,2-glucans and comparison with known SGLs.

| Enzyme | $K_m$ (mM) | $k_{cat}$ ($s^{-1}$) | $k_{cat}/K_m$ ($s^{-1} mM^{-1}$) |
| --- | --- | --- | --- |
| XccOpgD | $0.028 \pm 0.0035$ | $0.34 \pm 0.015$ | $12 \pm 1.1$ |
| EcOpgD<sup>a</sup> (GH186) | $0.44 \pm 0.056$ | $29 \pm 2$ | $67 \pm 3.5$ |
| CpSGL<sup>a</sup> (GH144) | $0.068 \pm 0.014$ | $49 \pm 4$ | $720 \pm 110$ |
| TfSGL<sup>a</sup> (GH162) | $0.015 \pm 0.001$ | $31 \pm 1$ | $2100 \pm 200$ |

<sup>a</sup>The values for EcOpgD, CpSGL and TfSGL are cited from Motouchi et al.<sup>12</sup>, Abe et al.<sup>14</sup> and Tanaka et al.<sup>15</sup>, respectively.

### Enzymatic properties of XccOpgD

XccOpgD exhibited the highest activity at 20–30 °C and pH 7.5 when linear β-1,2-glucans were used as substrates (Fig. 2a, b). XccOpgD exhibited no activity toward various polysaccharides, except linear β-1,2-glucan, indicating that XccOpgD is highly specific toward linear β-1,2-glucans (Figs. 1a, 2c). Thus, kinetic analysis of XccOpgD was performed using linear β-1,2-glucan as the substrate (Fig. 2d). A quantitative method for the cyclized product developed in this study is illustrated in Fig. 2e (see Methods for details). Although the *k*_cat_ value of XccOpgD was slightly lower than expected for a GH enzyme, the *K*_m_ value of XccOpgD was much lower than those of EcOpgD (GH186) and comparable to other β-1,2-glucan-related GHs, i.e., SGL from *Chitinophaga pinensis* and SGL from *Talaromyces funiculosus* (GH144 and GH162, respectively)^12,14,15^, leading to a *k*_cat_/*K*_m_ indicative of GH enzymes^16^ (Table 1).

**Fig. 2.**
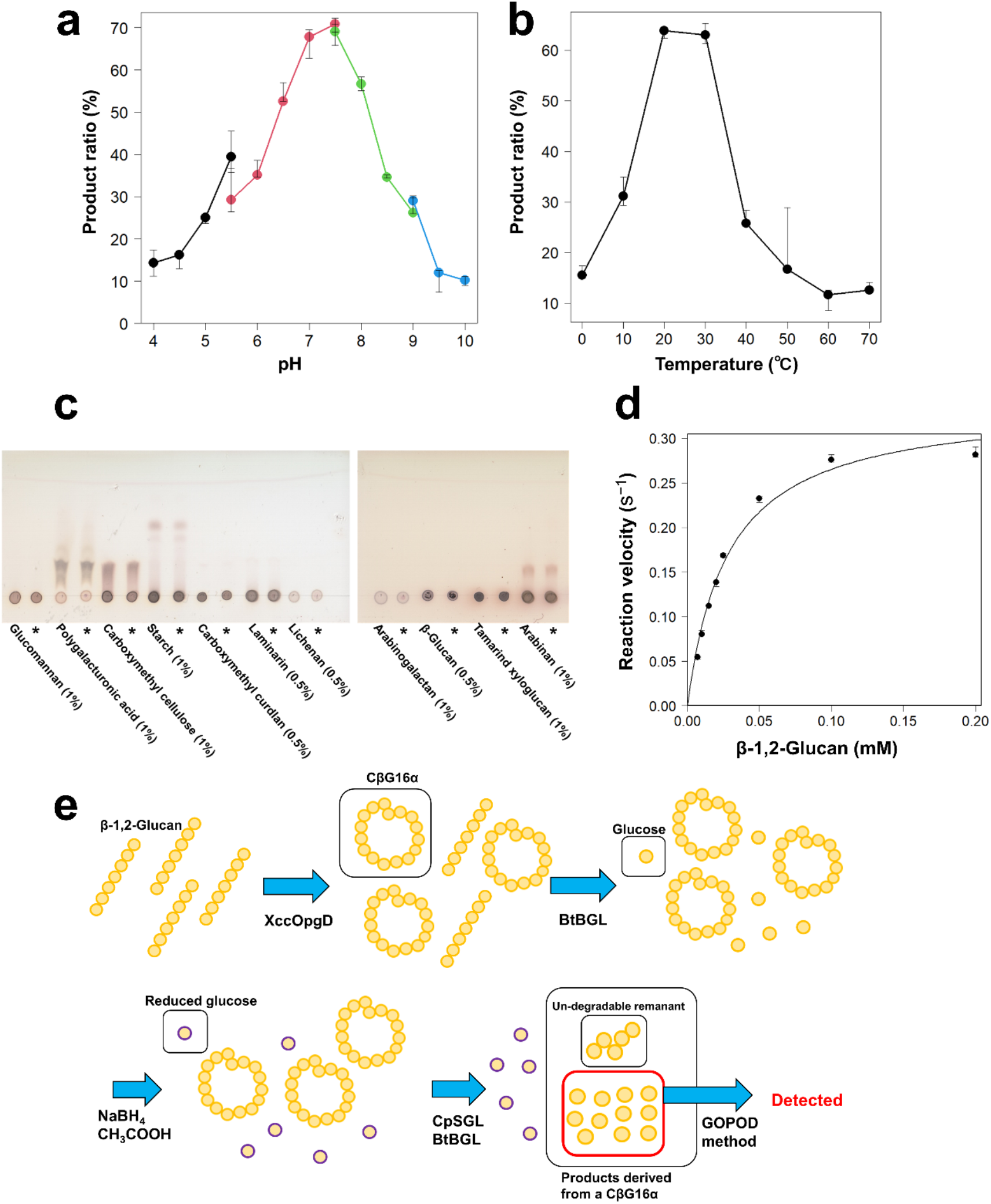
Enzymatic properties of XccOpgD. **a**, pH optimum. Buffers used for the enzymatic reactions were sodium acetate-HCl (pH 4.0–5.5, black), bis-Tris-HCl (pH 5.5–7.5, red), Tris-HCl (pH 7.5–9.0, green) and glycine-NaOH (pH 9.0–10, blue). **b**, Temperature optimum. **c**, Substrate specificity of XccOpgD. The reaction was performed at 37 °C for 24 h. Asterisks indicate that the reaction time was 24 h. The other lanes represent a reaction time of 0 h. **d**, Kinetic analysis of XccOpgD. Data plotted as closed circles are medians from triplicate experiments, and the other data were used for error bars (**a**, **b**, **d**). Data were regressed with the Michaelis-Menten equation (solid line). **e**, Strategy for quantifying the specific activity of XccOpgD.

### Michaelis complex of XccOpgD

The D379N mutant was used to determine a Michaelis complex structure of XccOpgD because D379 is conserved in GH186 and equivalent to the general acid of EcOpgD (Extended Data Fig. 3)^12^. The complex structure was obtained as a dimer (Fig. 3a). Three glucan chains were observed in this structure. Two glucans have DPs of 11 and 10 and bind to sites distal from the catalytic pocket of XccOpgD (Fig. 3a,b).

**Fig. 3.**
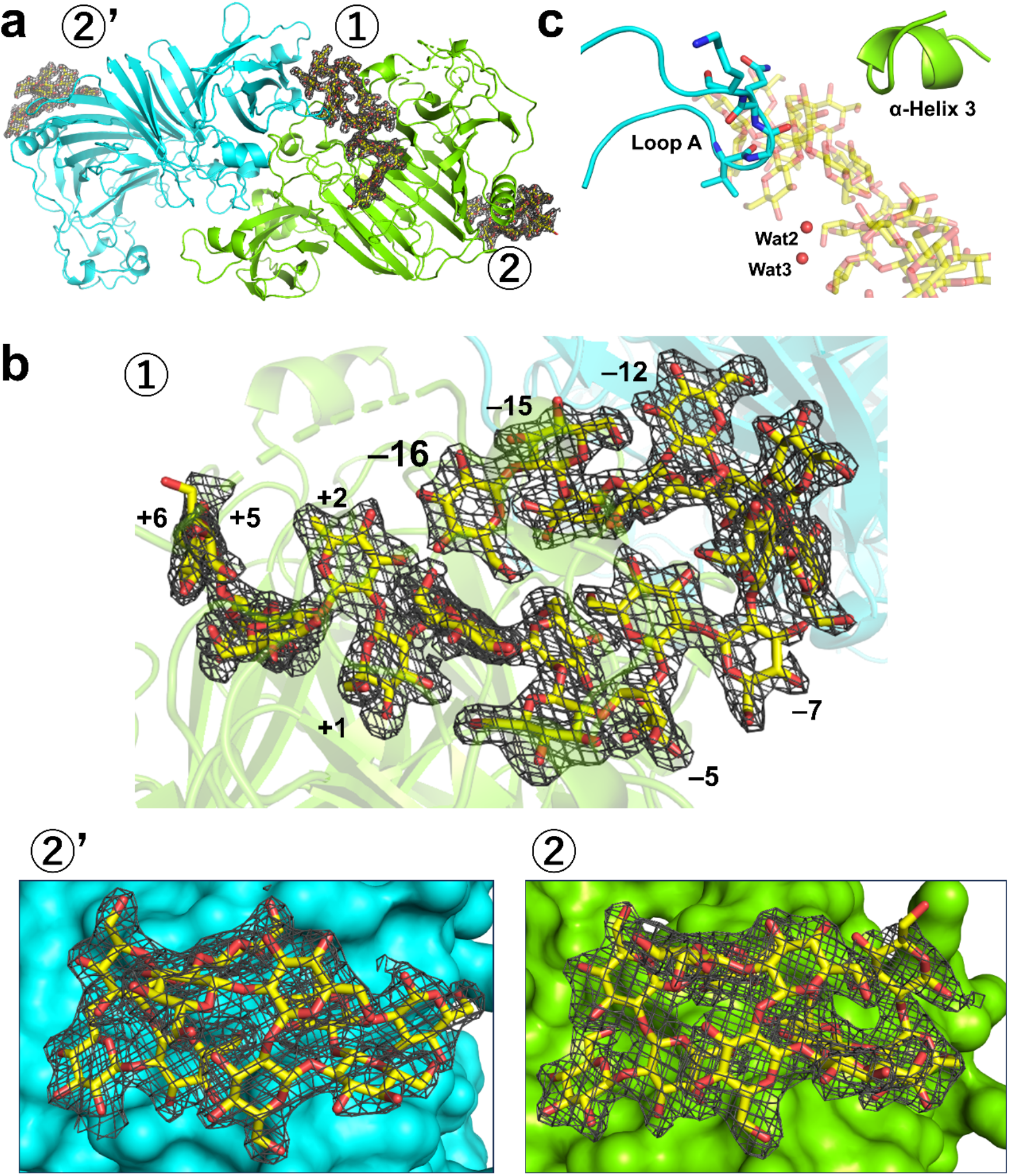
Structure of the Michaelis complex of XccOpgD. Chains A and B are shown in light green and cyan, respectively. Linear β-1,2-glucans are shown as yellow sticks. **a**, An asymmetric unit of XccOpgD and observed linear β-1,2-glucans. **b**, Close-up view around each linear β-1,2-glucan. The electron densities of linear β-1,2-glucans are shown as the *F*_o_–*F*_c_ omit maps by grey meshes at the 3σ contour level. **c**, Overview around Loop A and α-Helix 3. Wat2 and Wat3 are shown as red spheres. Linear β-1,2-glucan is shown semi-translucently.

In the catalytic pockets of the dimer, the electron density of a linear β-1,2-glucan with DP22 was clearly observed only in Chain A (Fig. 3b, top), probably because the closure motion of the cleft around α-Helix 3 is inhibited by crystal packing in Chain B (Fig. 3c, Extended Data Fig. 3), implying that the closure motion of the region is needed for substrate recognition. Superposition between Michaelis complexes of EcOpgD (PDB ID: 8IP1) and XccOpgD indicated that Chains A and B of the XccOpgD Michaelis complex form closed and open states, respectively (Extended Data Fig. 4). Thus, Chain A was used for describing the complex.

### Substrate binding mode of XccOpgD

Superimposed with the EcOpgD Michaelis complex, the positions and conformations of Glc moieties at subsites –7 to +6 (subsite is the nomenclature used for substrate binding sites, see https://www.cazypedia.org/index.php/Sub-site_nomenclature for details^17^) in XccOpgD are well conserved (Fig. 4a). In particular, the distorted conformation (^1^*S*_3_) of the Glc moiety at subsite –1 is well superimposed (Fig. 4b, Extended Data Fig. 5). The subsite positions of XccOpgD are the same as EcOpgD after considering that such distortion generally causes energetic instability for transition state in an enzymatic reaction (Fig. 4a). In XccOpgD, the substrate is also observed from subsites –16 to –8 (Fig. 4c). Subsite –16 is located near subsite –1, which appears to be suitable for cyclization (Fig. 3b). This is consistent with the compound produced by XccOpgD being α-1,6-cyclized β-1,2-glucohexadecaose.

**Fig. 4.**
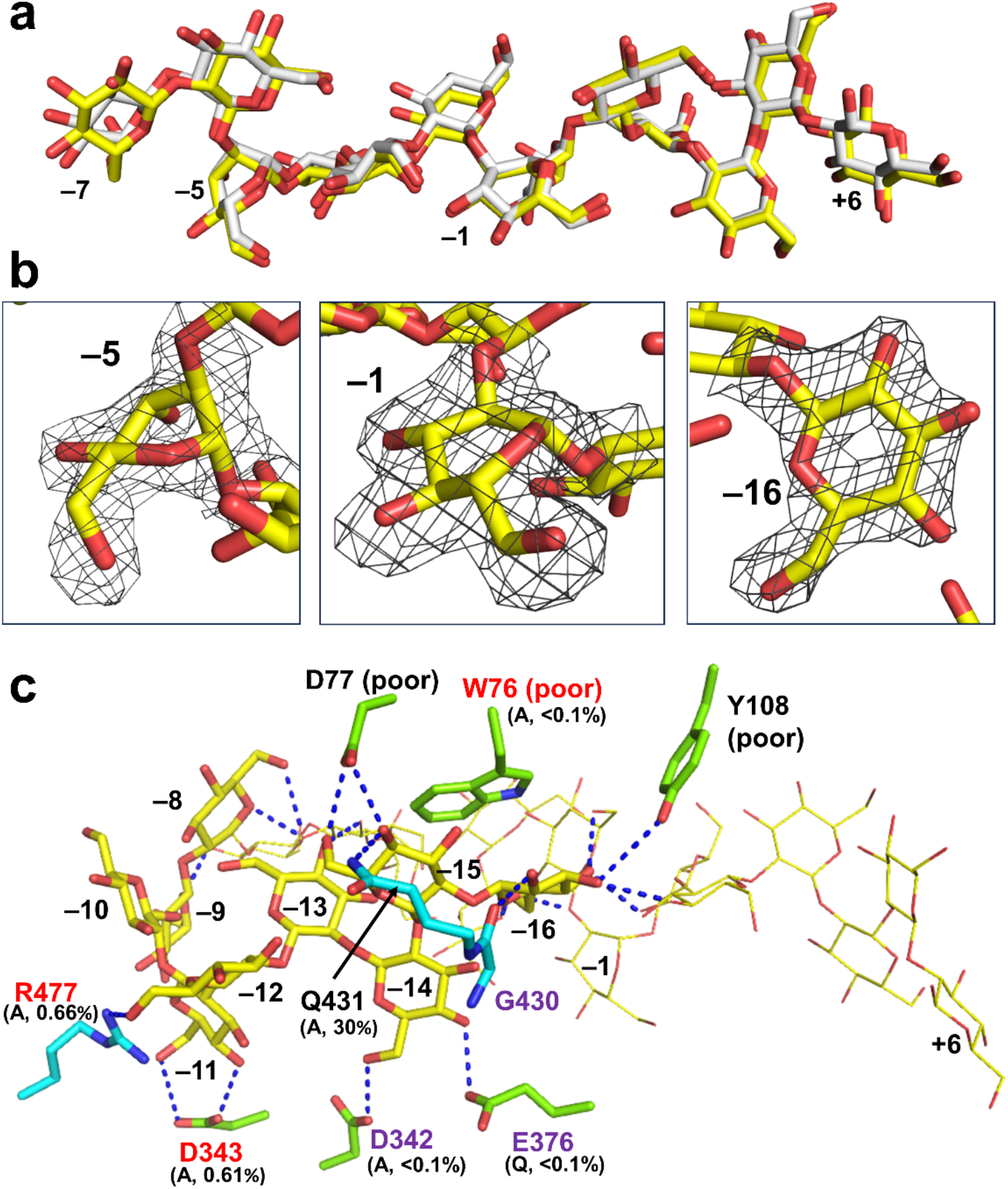
Substrate binding mode of XccOpgD. Linear β-1,2-glucans of XccOpgD and EcOpgD are shown as yellow and white sticks, respectively, except in part. **a**, Superimposition of linear β-1,2-glucans of XccOpgD and EcOpgD at subsites –7 to +6. **b**, The conformations of Glc moieties at subsites –5, –1 and –16 of XccOpgD. The electron density of each Glc moiety is shown as the *F*_o_–*F*_c_ omit map by grey mesh at the 3σ contour level. **c**, Substrate binding mode at subsites –16 to –8 of XccOpgD. The Glc moieties at subsites –7 to +6 are shown in yellow line representation. Residues in Chains A and B are shown as light green and cyan sticks, respectively. The relative activities (%) of mutants are shown below the substituted residues in parentheses with substituting residues. The residues highly conserved in GH186 are shown in purple letters. Residues whose mutations caused loss of cyclization activity despite not being conserved in GH186 are shown in red letters. Residues with relatively poor electron densities are indicated as (poor).

In EcOpgD, an α-helix moves drastically to form the catalytic cleft tunnel; however, P68 and N72 in α-Helix 3, which bind a substrate in the Michaelis complex, are not important for catalytic efficiency^12^. In contrast, in XccOpgD W76 in α-Helix 3 is likely to be important because this residue forms a stacking interaction with the Glc moiety at subsite –16 (Fig. 3c, 4c), and this is consistent with the W76A mutant displaying no cyclization activity (<0.1% specific activity compared with wild-type XccOpgD) (Table 2). In addition, CD spectra of all mutants gave similar spectra (Supplementary Fig. 1). The Glc moiety at subsite –5 in EcOpgD is distorted (Fig. 4b); however, the reason why such distortion is required is unclear^12^. Interestingly, such distortion is conserved in XccOpgD, which is important for orientating a linear β-1,2-glucan to achieve a cyclic form (Fig. 4).

**Table 2.** Specific activities of XccOpgD mutants.

|  | Specific activity (U/mg) | Relative activity (%) |
| --- | --- | --- |
| D379N | ND | ND |
| D291N | ND | ND |
| R350A | ND | ND |
| E376Q | ND | ND |
| Y347F | 0.012 (0.00081) | 4.2 |
| T377S | 0.016 (0.00030) | 5.5 |
| W76A | ND | ND |
| Q431A | 0.086 (0.0059) | 30 |
| R477A | 0.0019 (0.00025) | 0.66 |
| D342A | ND | ND |
| D343A | 0.0018 ($9.8 \times 10^{-5}$) | 0.61 |
| Wild-type | 0.29 | 100 |

All specific activities were measured using 0.1 mM linear β-1,2-glucan as the substrate.
Medians of triplicate experiments are presented.
ND represents data of less than 0.00029 units/mg (0.1% relative activity)
Maximum absolute values between medians and the other data of the triplicate experiments are shown in parentheses.
Activity to produce 1 $\mu$ mol of reaction product per minute is defined as 1 U ( $\mu$ mol/min).

There are relatively few substrate recognition residues at subsites –16 to –8 than at subsites –7 to +6 (Fig. 4c, Extended Data Fig. 3). Only intramolecular hydrogen bonds are found at four out of nine of these subsites (i.e., –8, –9, –10 and –13), which compensates for the stability of the binding mode (Fig. 4c, Extended Data Fig. 6). Among the substrate recognition residues for subsites –16 to –11, W76, D342, D343, E376, Q431 and R477 appear to be crucial (Fig. 4c). W76A, D342A, D343A, E376Q and R477A mutations severely disrupted cyclization activity (Table 2, Fig. 4c). Among these mutated residues, D342 and E376 are highly conserved in GH186. The other residues are conserved only in limited clades, indicating that residues corresponding to W76, D343 and R477 may distinguish α-1,6-cyclases from other GH186 enzymes (see Discussion for further detail).

### Unprecedented transglycosylation mechanism of XccOpgD

Anomer-inverting hydrolysis is the reaction where the anomeric orientation at the scissile bond is inverted after hydrolysis. EcOpgD adopts an anomer-inverting hydrolytic mechanism, and the general acid and base are likely to be D388 and D300, respectively^12^. A unique feature of the reaction mechanism is that D300 requires a route via a 4-hydroxy group of the Glc moiety at subsite –1 and two water molecules (Wat3 and Wat2) to deprotonate nucleophilic water (Wat1) by proton transfer (Extended Data Fig. 5)^12^.

To understand the reaction mechanism of XccOpgD, we compared XccOpgD with EcOpgD in terms of three key points: general acid, nucleophile and general base. The general acid of EcOpgD (D388), located at a suitable position for protonating a glycoside bond between subsites –1 and +1, is conserved in XccOpgD as D379. The Glc moiety at subsite –1 required for hydrolytic activity is distorted in XccOpgD, as observed for EcOpgD (Fig. 5b). Thus, D379 is probably the general acid, and the dramatic loss of cyclization activity (<0.1%) of the D379N mutant supports this residue acting as a general acid (Table 2).

**Fig. 5.**
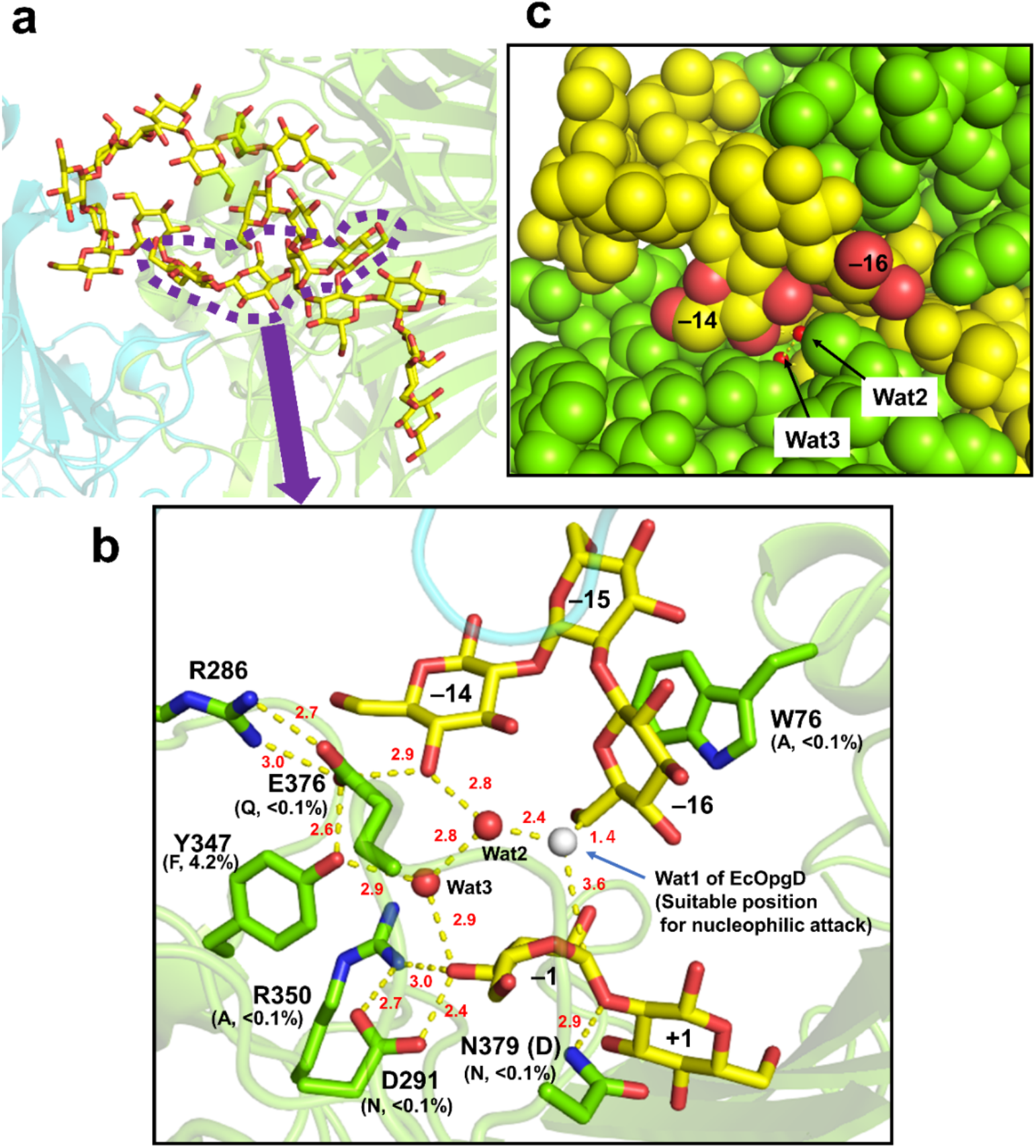
Environment around a linear β-1,2-glucan molecule bound to the catalytic cleft of XccOpgD. Hydrogen bonds are indicated by yellow dotted lines with lengths (red, Å). Residues and substrates are shown as green and yellow sticks, respectively. **a**, Conformation of the whole linear β-1,2-glucan molecule in XccOpgD. Chains A and B are shown as semi-transparent green and cyan cartoons, respectively. **b**, Close-up view around subsites –14 and –16. The structure is superimposed with the Michaelis complex of EcOpgD (PDB ID: 8IP1) to show Wat1, a nucleophilic water (white sphere) of EcOpgD, as a suitable position for nucleophilic attack. The relative activities (%) of mutants are shown below the substituted residues in parentheses with substituting residues. (D) by N379 represents the original residue in wild-type XccOpgD. **c**, The sequestered proton transfer pathway of XccOpgD. Chain A and the substrate are shown as light green and yellow spheres, respectively. Wat2, Wat3 and oxygen atoms of the Glc moieties at subsites –14 and –16 are shown as red spheres. Spheres except for Wat2 and Wat3 are shown with van der Waals radii.

Nevertheless, unlike EcOpgD, a nucleophilic water molecule is absent in XccOpgD. In XccOpgD, the 6-hydroxy group of the Glc moiety at subsite –16 is located near the anomeric carbon of the Glc moiety at subsite –1 (Fig. 5b). Although the 6-hydroxy group does not adopt a suitable orientation for nucleophilic attack, the distance between Wat1 in EcOpgD and C6 of the Glc moiety at subsite –16 is 1.4 Å, which is a covalent bond distance (Fig. 5b). This observation suggests that the 6-hydroxy group of the Glc moiety at subsite –16 can locate to a suitable position for nucleophilic attack. Therefore, anomer-inverting transglycosylation will occur if the 6-hydroxy group is deprotonated.

Subsequently, to identify a general base, we traced the proton transfer pathway from Wat1 of EcOpgD, a potential position of the 6-hydroxy group of the Glc moiety at subsite –16. The pathway of Wat2-Wat3-O4 (subsite –1)-D291 of XccOpgD is the same as EcOpgD and the D291N mutant lost cyclization activity (<0.1%) (Fig. 5b, Extended Data Fig. 5, Table 2), suggesting that D291 is a strong candidate for a general base. Nevertheless, E376 may also be a general base candidate (Fig. 5b). The pathway of Wat2-Wat3-Y347-E376 is probably not the catalytic pathway because the Y347F mutant did not fully lose cyclization activity (4.2%) (Fig. 5b, Table 2). The other pathway to reach E376, Wat2-O4 (subsite −14)-E376, may be possible because the E376Q mutant lost cyclization activity (<0.1%) (Fig. 5b, Table 2). However, the remarkable decrease in cyclization activity for the E376Q mutant is probably because E376 is a substrate recognition residue at minus subsites near the non-reducing end, as described above. Indeed, D342A, which is the same mutant as E376Q in terms of losing one substrate recognition at subsite –14, showed minimal levels of cyclization activity (<0.1%) (Table 2, Fig. 4c). Therefore, we propose that D291 is the general base, whereas E376 is probably only important for substrate recognition.

In EcOpgD, the unprecedentedly long proton transfer from Wat1 was suggested to occur efficiently because Loop A (residues 434–453 a.a.) sequesters the proton transfer pathway from solvent^12^ (Extended Data Fig. 3). Although Loop A in XccOpgD is too short to cover the proton transfer pathway, the proton transfer pathway is sequestered from solvent by the Glc moieties at subsites –14 and –16 (Fig. 5c), which accounts for the efficient proton transfer of XccOpgD. Overall, the reaction mechanism proposed in Fig. 6 is the first-discovered anomer-inverting transglycosylation.

**Fig. 6.**
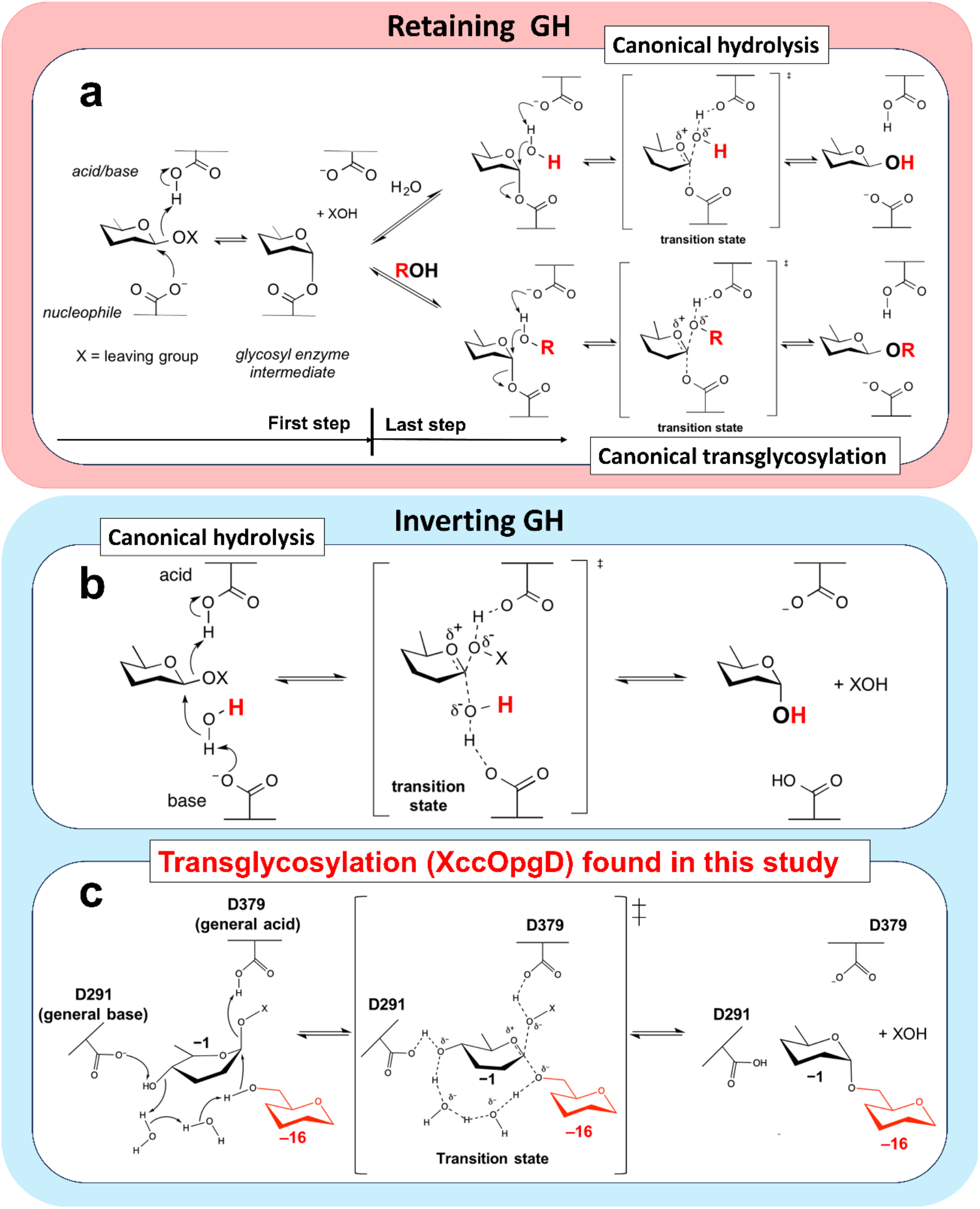
A proposed anomer-inverting transglycosylation mechanism of XccOpgD and comparison with canonical hydrolysis and transglycosylation. Arrows represent the pathway for electron transfer. The groups transferred to the anomeric position are shown as red letters or red structural formula. **a,b**, The figures are cited by CAZypedia^19^ (https://www.cazypedia.org/index.php/) and partly modified. **a–c**, The reaction mechanism of anomer-retaining hydrolysis and transglycosylation (**a**), anomer-inverting hydrolysis (**b**) and transglycosylation (**c**).

## Discussion

α-1,6-Cyclized β-1,2-glucan is produced by *X. campestris*, a well-known plant pathogen, and by model photosynthetic bacteria such as *Rhodobacter sphaeroides*^2,4,18^. Interestingly, CβG16α was reported to be an immune-avoidance and infectious factor toward plants^3^. However, no enzyme that synthesizes α-1,6-cyclized β-1,2-glucan has been identified. In this study, we focused on XccOpgD based on a functional diversity perspective in the GH186 family and identified XccOpgD as the enzyme responsible for completing CβG16α synthesis. This enzyme should be given a new EC number. We propose linear (1→2)-β-D-glucan:(1→2)-β-D-glucohexadecaose 6-α-D-[(1→2)-β-D-glucohexadecaosyl]-transferase (cyclizing) as a systematic name and α-1,6-cyclized β-1,2-glucohexadecaose synthase as an accepted name.

Furthermore, structural and functional analyses of XccOpgD revealed an unprecedented anomer-inverting transglycosylation mechanism (Fig. 6c). Among glycoside hydrolases, reactions are categorized into two canonical types, anomer-retaining GH and anomer-inverting GH, based on CAZypedia (https://www.cazypedia.org/index.php/) (Fig. 6). In both types, hydrolysis occurs when a nucleophile in the last (second) step of the reaction is water (Fig. 6a,b). In contrast, transglycosylation has long been considered to occur only as an anomer-retaining mechanism^19^ (Fig. 6a). Therefore, the XccOpgD reaction (Fig. 6c) is the first discovery of an anomer-inverting transglycosylation. Because the ratio of anomer-inverting GHs is approximately 40% in all reaction mechanism-determined GH families, our discovery sheds light on the long-held possibility of generating new glycan synthases by modifying anomer-inverting GHs.

This study corroborated the diversity of the reaction mechanisms in the GH186 family and provided perspectives on functionally unknown clades in GH186. Mutational analyses revealed that W76, D343 and R477 are important substrate recognition residues for the cyclization activity of XccOpgD to produce CβG16α. These residues represent indicators for distinguishing α-1,6-cyclized β-1,2-glucohexadecaose synthases from other GH186 homologues. In GH186, these three residues are highly conserved in only clade 3, which is next to the clade of EcOpgD in the phylogenetic tree (Extended Data Fig. 7a). Although W76 of XccOpgD is substituted with Tyr in some clade 3 homologues, the stacking interactions with the Glc moiety at subsite –16 are expected to be conserved, indicating that clade 3 homologues share efficient cyclization activity (Extended Data Fig. 7c).

In clade 4, W76 of XccOpgD is substituted with Tyr (Extended Data Fig. 7d), as observed for some homologues in clade 3, implying that homologues in clade 4 retain anomer-inverting transglycosylation activity. However, D343 and R477, important residues in XccOpgD, are not conserved in clade 4 (Extended Data Fig. 7d). These features suggest that homologues in clade 4 produce glucose polymers with different DPs from CβG16α. Indeed, it was reported that *Rhodobacter sphaeroides* possessing the OpgD homologue in clade 4 synthesizes α-1,6-cyclized β-1,2-glucan with a DP of 18 (CβG18α)^18^.

In clade 2, particular homologues near clade 3 have conserved residues corresponding to R477 and D343 of XccOpgD, implying that they recognize the Glc moieties at minus subsites, which are important for locating linear β-1,2-glucan as a suitable form for cyclization (Extended Data Fig. 7b). Nevertheless, these homologues have the highly conserved GW motif on Loop A, which is vital for stabilizing the nucleophilic water required for hydrolytic activity^12^. Furthermore, W76 of XccOpgD is a proline, which cannot form stacking interactions with a Glc moiety, suggesting that these homologues are not transglycosylases but hydrolases. Thus, it is likely that the homologues described above are intermediates of EcOpgD and XccOpgD and that the significant changes in the reaction mechanisms arose from the minor sequence variations described above.

Overall, GH186 homologues that produce α-1,6-cyclized β-1,2-glucohexadecaose are hypothesized to be limited to clade 3, and the function of many GH186 homologues differs from XccOpgD and EcOpgD. Homologues of most *Xanthomonas* pathogens are located specifically in clade 3, implying that many *Xanthomonas* diseases can be regulated by inhibiting α-1,6-cyclized β-1,2-glucohexadecaose synthase. Furthermore, given that the environment for capturing subsites –7 to +6 is highly conserved in the majority of GH186 enzymes, GH186 can be a broadly adaptable inhibition target for treating *Xanthomonas* diseases related to CβG16α and for other animal or plant pathogens that use other OPGs such as LβG-6β.

The discovery of an unprecedented glycan-generating mechanism in this study provides new possibilities for the enzymatic synthesis of carbohydrates. Furthermore, identifying enzymes that synthesize CβG16α is a future milestone for understanding plant infection systems. GH186 enzymes will offer valuable inhibition targets for creating new-genre agrichemicals that have the potential to answer one of the biggest agrichemical issues, antimicrobial-resistant bacteria.

## Methods

### Cloning and purification of XccOpgD

The gene encoding XccOpgD (GenBank: AAM43366.1) was amplified by PCR with the primer pair shown in Supplementary Table 1 using PrimeSTAR Max (Takara Bio) and a genomic DNA of *X. campestris* pv. *campestris* (DSM3568, the Leibniz Institute, Germany) as the template. The forward primer was designed to eliminate the N-terminal signal sequence predicted by the SignalP6.0 server (https://services.healthtech.dtu.dk/services/SignalP-6.0/)^20^. The amplified gene was inserted between the XhoI and NdeI sites of the pET30a vector by the SLiCE method^21^ to add a C-terminal His_6_-tag to the target protein. The constructed plasmid was transformed into *E. coli* Rosseta2 (DE3) cells and the transformants were cultured in 1 L Luria-Bertani medium containing 30 mg/L kanamycin at 37 °C until the absorbance at 600 nm reached 0.6. Expression was induced by adding isopropyl β-D-1-thiogalactopyranoside to a final concentration of 0.1 mM, and cells were cultured at 20 °C for 24 h. The cells were centrifuged at 7000 *g* for 10 min and resuspended in 50 mM Tris-HCl buffer (pH 7.5). The resuspended cells were disrupted by sonication, and the sample was centrifuged at 33000 *g* for 15 min to obtain a cell extract. The cell extract was loaded onto a HisTrap™ FF crude column (5 mL; Cytiva) pre-equilibrated with buffer (50 mM Tris-HCl, 500 mM NaCl and 20 mM imidazole, pH 7.5). After washing the column with the same buffer, the target protein was eluted with a linear gradient of 20–300 mM imidazole in a buffer containing 50 mM Tris-HCl (pH 7.5) and 500 mM NaCl. Amicon Ultra 30,000 molecular weight cutoff centrifugal filters (Merck, NJ, USA) were used to concentrate a portion of the fractionated protein and exchange the buffer to 50 mM Tris-HCl (pH 7.5) containing 50 mM NaCl. Each purified protein migrated as a single band of 60 kDa on SDS-PAGE gels, which is consistent with the theoretical molecular mass of XccOpgD (57851.479 Da). Concentrations of purified enzymes were calculated from the absorbance at 280 nm^22^. The extinction coefficient of XccOpgD at 280 nm is 115270 mol^−1^cm^−1^.

### β-1,2-Glucans used for experiments

Linear β-1,2-glucans with an average DP of 121 calculated from their number average molecular weight (Mn) were used for TLC analysis, product preparation for ESI-MS and NMR, investigation of general properties and kinetic analysis. The average DP of linear β-1,2-glucans used for crystallization to obtain the Michaelis complex of XccOpgD was 17.7 based on the Mn. All linear β-1,2-glucans used in this study were prepared according to several references^23,24^.

### TLC analysis

XccOpgD (0.55 mg/mL) was incubated with 1% linear β-1,2-glucan in 2.5 mM Tris-HCl buffer (pH 7.5) at 37 °C for 24 h (sample 1). One microliter 1 mg/mL BtBGL was added to 10 μL sample 1 and incubated at 37 °C for 24 h (sample 2). One microliter 1 mg/mL BtBGL and 1 μL 3 mg/mL CpSGL were added to sample 1 and incubated at 37 °C for 24 h (sample 3). The reaction mixtures (1 μL) were spotted onto TLC Silica Gel 60 F_254_ (Merck) plates. The plates were developed with 72% acetonitrile once. The plates were then soaked in a 5% (w/v) sulfuric acid/methanol solution and heated in an oven until the spots were clearly visualized. A mixture of β-1,2-glucooligosaccharides was prepared as a marker by incubating β-1,2-glucooligosaccharides with DPs 3–7 and linear β-1,2-glucan in 1 mM sodium phosphate containing 1,2-β-oligoglucan phosphorylase from *Listeria innocua*^25^.

### ESI-MS

XccOpgD (0.135 mg/mL) was incubated with 1% linear β-1,2-glucan in 5 mM Tris-HCl buffer (pH 7.5) at 37 °C for 24 h (sample 1). One microliter 10 mg/mL BtBGL was added to 49 μL sample 1 and incubated at 37 °C for 24 h (sample 2). 0.24 mg/mL BtBGL and 0.060 mg/mL CpSGL were incubated with 1% purified CβG16α at 37 °C for 24 h (sample 3). Amberlite MB4 (Organo) was added to samples 1 and 2 to remove ionic compounds. The resultant solutions were diluted 100-fold with a solvent (methanol/water = 1/1, v/v) containing 5 mM ammonium acetate. After filtration, the samples were loaded onto the Sciex X500 R QTOF (Sciex) in the positive mode at a 20 μL/min flow rate.

### NMR

The enzymatic reaction of 20 μg/mL XccOpgD, 1% linear β-1,2-glucan and 4 mM Tris-HCl buffer (pH 7.5) was incubated at 30°C overnight. Then, 10 μL 109 mg/mL BtBGL and 250 μL 1M bis-Tris buffer (pH 5.5) was added and incubated overnight. The reaction product was purified by size-exclusion chromatography using a Toyopearl HW-40F column (∼2 L gel). The sample was eluted with distilled water after the injection of the reaction mixture (∼10 mL). The eluates were fractionated into 30-mL portions, and the fraction containing only the main product of the reaction by XccOpgD was lyophilized. One-(^1^H and ^13^C) and two-dimensional (double-quantum-filtered COSY, heteronuclear single-quantum coherence (HSQC), heteronuclear multiple-bond correlation (HMBC) and HSQC-TOCSY) NMR spectra of the products were acquired in D_2_O at 298 K using a Bruker Avance 800 MHz spectrometer (Bruker, MA, USA) with *t*BuOH (*δ* 1.23 ppm for ^1^H, and *δ* 31.30 ppm for ^13^C) as an internal standard. The H-1 chemical shift at the α-anomer was assigned by referring to the Karplus curve^26^. The H-2 chemical shift was assigned based on double-quantum-filtered COSY spectra. The C-1 and C-2 chemical shifts were assigned using HSQC spectra and based on the assignment of H-1 and H-2 proton signals. The H-6 and C-6 chemical shifts were determined using DEPT135, HSQC and HSQC-TOCSY data. Linkage positions between glucose units were determined by detecting cross-peaks in the HMBC spectrum.

### General properties

The optimum pH of XccOpgD (0.1 mg/mL) activity was determined by incubating the protein in various 20 mM buffers (sodium acetate-HCl, pH 4.0–5.5; bis-Tris-HCl, pH 5.5–7.5; Tris-HCl, pH 7.5– 9.0; glycine, pH 9.0–10.0) containing 0.25% linear β-1,2-glucan at 30 °C for 10 min and then heating at 100 °C for 5 min to terminate the reaction. BtBGL and bis-Tris-HCl (pH 5.5) were added (final concentrations were 1 mg/mL and 83 mM, respectively) to convert non-cyclized β-1,2-glucan into glucose. The amounts of produced glucose were measured by the GOPOD method^16^ to determine the amounts of residual linear β-1,2-glucan. The conversion rates were calculated as subtracts between weights of the initial and residual substrates in the reaction by XccOpgD. The optimum temperature was determined by performing the reactions in 20 mM Tris-HCl buffer (pH 7.5) at each temperature (0–70 °C) for 10 min and then heated at 100 °C for 5 min to terminate the reaction. The conversion rates were calculated using the same approach to determine the optimum pH.

### Substrate specificity

XccOpgD (0.61 mg/mL) was incubated in 5 mM Tris-HCl buffer (pH 7.5) containing each substrate (1% glucomannan, Neogen, MI, USA; 1% polygalacturonic acid, Neogen; 1% carboxymethyl cellulose, Merck; 1% soluble starch, FUJIFILM Wako Chemical Corporation, Osaka, Japan; 0.5% carboxymethyl curdlan, Neogen; 0.5% laminarin, Merck; 0.5% lichenan, Neogen; 1% arabinogalactan, Neogen; 0.5% barley β-glucan, Neogen; 1% tamarind-xyloglucan, Neogen; or 1% arabinan, Neogen) at 30 °C for 24 h. The reaction patterns were analysed by TLC.

### Kinetic analysis

The kinetic parameters for linear β-1,2-glucans were determined by performing the enzymatic reaction in a 20 μL reaction mixture containing 0.2 mg/mL XccOpgD, 0.007–0.2 mM linear β-1,2-glucan and 20 mM Tris-HCl (pH 7.5) at 30 °C for 10 min. The reaction was stopped by heat treatment at 100 °C for 5 min. BtBGL and bis-Tris-HCl (pH 5.5) were added (final concentrations 1 mg/mL and 16.7 mM, respectively) to convert non-cyclized β-1,2-glucans into glucose. The reaction products were reduced using a one-fifth volume of 1 M NaBH_4_. The same volume of 1 M acetate as that of the 1M NaBH_4_ solution was added to each sample to neutralize NaBH_4_-treated solutions. The samples were then treated with 0.6 mg/mL of BtBGL and 0.15 mg/mL CpSGL at 30 °C for 24 h to convert CβG16α into 11 glucoses and a non-degradable glucopentaose. The main residual oligosaccharide was identified to be glucopentaose by ESI-MS.

Colour development of the reaction mixtures was performed using the GOPOD method^16^ to quantify the concentration of glucose derived from the products released by XccOpgD. Molar concentrations of linear β-1,2-glucan substrates were calculated based on the Mn of the substrates. Kinetic parameters of XccOpgD were determined by fitting experimental data to the Michaelis-Menten equation, *v*/[E]_0_ = *k*_cat_ [S]/(*K*_m_+ [S]), where *v* is the initial velocity, [E]_0_ is the enzyme concentration, [S] is the substrate concentration, *k*_cat_ is the turnover number, and *K*_m_ is the Michaelis constant. Each analysis was performed in triplicate, and the medians were used for regression.

### Crystallography

As described above, XccOpgD (D379N) was purified using a HisTrap™ FF crude column. The crystal of the D379N mutant for the linear β-1,2-glucan complex was obtained at 20 °C after a month by mixing 1 μL D379N mutant (7.0 mg/mL) with 1 μL reservoir solution comprising 0.1 M bis-Tris buffer (pH 5.5) and 2 M (NH_4_)_2_SO_4_. The complex crystal of D379N was soaked in the reservoir solution supplemented with 30% (w/v) glycerol and 20% (w/v) linear β-1,2-glucan. The crystal was kept at 100 K in a nitrogen-gas stream during data collection. The X-ray diffraction data was collected on a beamline (BL-5A) at Photon Factory (Tsukuba, Japan). The diffraction data of crystal of the linear β-1,2-glucan-bound XccOpgD (D379N) was collected at 1.0 Å and processed with X-ray Detector Software (http://xds.mpimf-heidelberg.mpg.de/)^27^ and the Aimless program (http://www.ccp4.ac.uk/). The initial phase of XccOpgD structure was determined by molecular replacement using the Alphafold2 predicted XccOpgD as a model structure. Molecular replacement, auto model building and refinement were performed using MOLREP, Buccaneer, REFMAC5 and Coot programs, respectively (http://www.ccp4.ac.uk/)^28–31^. Crystallographic data collection and refinement statistics are summarized in Extended Data Table 3. All visual representations of the structures were prepared using PyMOL (https://pymol.org/2/).

### Mutational analysis

The plasmids for producing XccOpgD mutants were constructed using a PrimeSTAR mutagenesis basal kit (Takara Bio) according to the manufacturer’s instructions. PCRs were performed using appropriate primer pairs (Supplementary Table 1) and the XccOpgD plasmid as the template. Transformation into *E. coli* Rosseta2 (DE3) and the expression and purification of XccOpgD mutants were performed using the same methods described for wild-type XccOpgD preparation. The enzymatic reactions of XccOpgD mutants were performed similarly to determine the specific activity at 0.1 mM substrate concentration. The final assay concentration of the mutants and reaction times were 0.023–2.6 mg/mL and 0–6.5 h, respectively, depending on the mutants. Colour development was performed in the same way as described in Kinetic analysis.

### CD spectra

CD spectra were recorded between 200–250 nm using a J820 spectropolarimeter (JASCO). Each sample contained 2 mM Tris-HCl (pH 7.5) and a mutant (0.011 mg/mL).

### Sequence analysis

All alignment was performed using Clustal Omega^32^ and visualized using the ESPript 3.0 server (http://espript.ibcp.fr/ESPript/ESPript/)^33^.

### Statistics and reproducibility

Quantitative data describing the activity of XccOpgD were obtained from three independent experiments, and medians were reported. The multiplicity of obtained structures is more than 10.

### Reporting summary

Further information on research design is available in the Nature Portfolio Reporting Summary linked to this article.

## Acknowledgements

This work was supported by Photon Factory for X-ray data collection (Proposal No. 2020G527) and in part by JST SPRING (Grant Number JPMJSP2151). We thank Edanz (https://jp.edanz.com/ac) for editing a draft of this manuscript.

## Author contributions

S.M., H.N. and M.N. conceived the project and designed the experiments. S.M. expressed and purified XccOpgD. S.M. and M.N. performed data collection and processed X-ray crystallographic data. K.S. performed NMR analyses. H.N. provided β-1,2-glucooligosaccharides. S.M., K.S. and M.N. prepared the manuscript. H.N. and M.N. supervised the project and participated in manuscript writing. All authors contributed to the revision of the manuscript. Any correspondence should be addressed to M.N. and S.M.

## Data availability

Atomic structure coordinates were deposited in the PDB under accession code 8X18. Original NMR data analysis is provided in Supplementary Data 1 (Supplementary_Data1.pdf)

## Extended Data for

**Extended Data Table 1.** H-1, H-2 and H-6 proton chemical shifts of the product produced by XccOpgD.

800 MHz NMR (D<sub>2</sub>O with internal standard tBuOH $\delta$ = 1.23, 298 K)
| Proton | Chemical shift of each glucose residue (ppm) |  |  |  |  |  |  |  |  |  |  |  |  |  |  |  |
| --- | --- | --- | --- | --- | --- | --- | --- | --- | --- | --- | --- | --- | --- | --- | --- | --- |
|  | 1 | 2 | 3 | 4 | 5 | 6 | 7 | 8 | 9 | 10 | 11 | 12 | 13 | 14 | 15 | 16 |
| H-1 (ppm) | 5.160 | 4.902 | 4.796 | 4.968 | 4.872 | 4.991 | 4.820 | 5.036 | 4.843 | 4.914 | 4.991 | 4.801 | 5.010 | 4.839 | 4.921 | 4.650 |
| $J_{1,2}$ (Hz) | 3.48 | 7.99 | 7.78 | 7.93 | 7.90 | 7.91 | 7.87 | 7.88 | 7.89 | 7.87 | 7.91 | 7.92 | 7.93 | 7.90 | 7.86 | 7.91 |
| H-2<sup>a</sup> (ppm) | 3.62 | 3.32 | 3.71 | 3.52 | 3.63 | 3.53 | 3.66 | 3.56 | 3.69 | 3.57 | 3.60 | 3.67 | 3.53 | 3.68 | 3.54 | 3.63 |
| H-6<sup>b</sup> (ppm) | A<sup>c</sup> | 3.85 | A<sup>c</sup> | A<sup>c</sup> | A<sup>c</sup> | A<sup>c</sup> | A<sup>c</sup> | A<sup>c</sup> | A<sup>c</sup> | A<sup>c</sup> | A<sup>c</sup> | A<sup>c</sup> | A<sup>c</sup> | A<sup>c</sup> | A<sup>c</sup> | A<sup>c</sup> |

<sup>a</sup>H-2 chemical shifts were determined using <sup>1</sup>H-<sup>1</sup>H COSY NMR data.
<sup>b</sup>H-6 chemical shifts were determined using DEPT135, HSQC, HSQC-TOCSY and HMBC NMR data.
<sup>c</sup>A: 3.73–3.98 ppm.

**Extended Data Table 2.** C-1, C-2 and C-6 carbon chemical shifts of the product produced by XccOpgD.

800 MHz NMR (D<sub>2</sub>O with internal standard tBuOH $\delta$ = 31.30, 298 K)
| Carbon | Chemical shift of each glucose residue (ppm) |  |  |  |  |  |  |  |  |  |  |  |  |  |  |  |
| --- | --- | --- | --- | --- | --- | --- | --- | --- | --- | --- | --- | --- | --- | --- | --- | --- |
|  | 1 | 2 | 3 | 4 | 5 | 6 | 7 | 8 | 9 | 10 | 11 | 12 | 13 | 14 | 15 | 16 |
| C-1 (ppm) | 100.02 | 104.03 | 104.73 | 103.28 | 103.94 | 102.85 | 103.65 | 102.80 | 103.91 | 103.46 | 102.69 | 104.16 | 102.98 | 104.18 | 103.94 | 104.80 |
| C-2 (ppm) | 83.55 | 75.26 | 82.33 | 85.96 | 83.36 | 84.63 | 81.62 | 84.63 | 81.05 | 84.22 | 83.31 | 81.38 | 85.04 | 81.38 | 85.06 | 83.69 |
| C-6 (ppm) | A<sup>a</sup> | 69.26 | A<sup>a</sup> | A<sup>a</sup> | A<sup>a</sup> | A<sup>a</sup> | A<sup>a</sup> | A<sup>a</sup> | A<sup>a</sup> | A<sup>a</sup> | A<sup>a</sup> | A<sup>a</sup> | A<sup>a</sup> | A<sup>a</sup> | A<sup>a</sup> | A<sup>a</sup> |

<sup>a</sup>A: 62.16–62.84 ppm.

**Extended Data Table 3.** Crystallographic data collection and refinement statistics of XccOpgD.

| Data set | XccOpgD-β-1,2-glucan<br>(D379N mutant) |
| --- | --- |
| **Data collection** |  |
| Beamline | KEK NW-12A |
| Space group | $P6_5$ |
| | $a = 171.88$ |
| | $b = 171.88$ |
| Unit cell parameters ($\text{\AA}$, $^\circ$) | $c = 128.69$ |
| | $\alpha, \beta = 90$ |
| | $\gamma = 120$ |
| Resolution ($\text{\AA}$)<sup>a</sup> | 48.723–2.25 (2.29–2.25) |
| Total reflections<sup>a</sup> | 1064039 (51345) |
| Unique reflections<sup>a</sup> | 102253 (5064) |
| Completeness (%)<sup>a</sup> | 100 (100) |
| Multiplicity<sup>a</sup> | 10.4 (10.1) |
| Mean $I/\sigma(I)$<sup>a</sup> | 11.6 (2.9) |
| $R_{\text{merge}}$ (%)<sup>a</sup> | 15.8 (85.6) |
| $R_{\text{pim}}$ (%)<sup>a</sup> | 7.6 (41.6) |
| $CC_{1/2}$<sup>a</sup> | (0.830) |
| **Refinement** |  |
| Resolution ($\text{\AA}$) | 48.723–2.25 |
| No. of reflections | 102207 |
| No. of atoms | 9012 |
| No. of water molecules | 541 |
| $R_{\text{work}}/R_{\text{free}}$ (%) | 17.3/17.6 |
| No. of asymmetric units | 2 |
| r.m.s.d. from ideal values |  |
| Bond lengths ($\text{\AA}$) | 0.0089 |
| Bond angles ($^\circ$) | 1.5083 |
| Average $B$-factors ($\text{\AA}^2$) | |
| Protein (chain A/B) | 31.1/34.8 |
| Ligand |  |
| β-1,2-Glucan | 39.11 |
| Solvent | 32.5 |
| Ramachandran plot (%) |  |
| Favoured | 97.0 |
| Allowed | 3.0 |
| Outlier | 0.0 |
| **PDB entry** | 8X18 |

<sup>a</sup>Values in parentheses represent the highest resolution shell.

**Extended Data Fig. 1.**
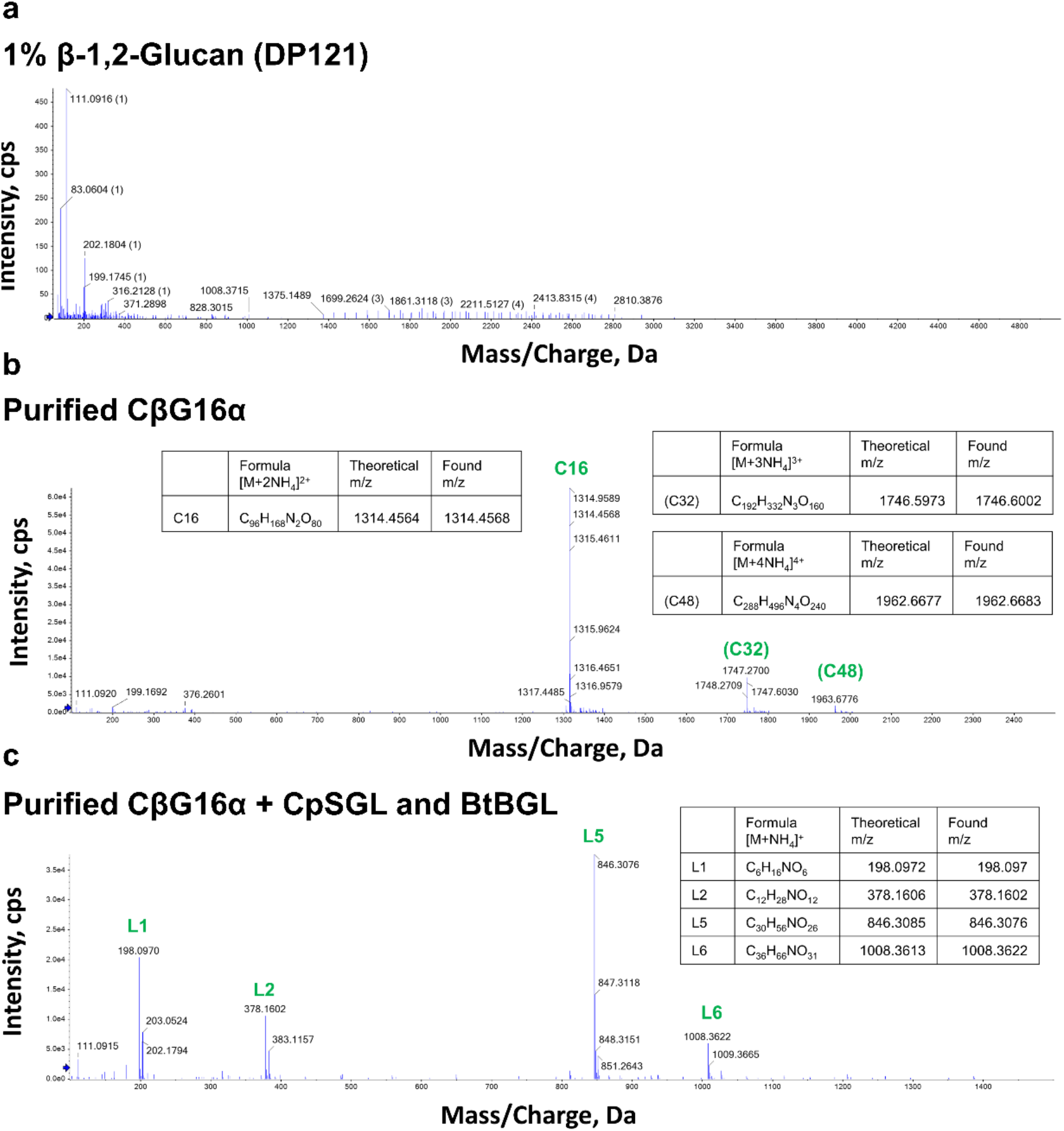
Electrospray ionization-mass spectrometry analysis. The peaks are assigned as [M + nNH_4_]^n+^ and indicated by arrows for the products. Green text represents forms of compounds (cyclic or linear) and DPs of products. For example, L5 represents a linear pentaose. **a–c**, ESI-MS data of linear β-1,2-glucan (**a**), purified CβG16α (**b**) and the reaction products released from CβG16α following treatment with BtBGL and CpSGL (**c**).

**Extended Data Fig. 2.**
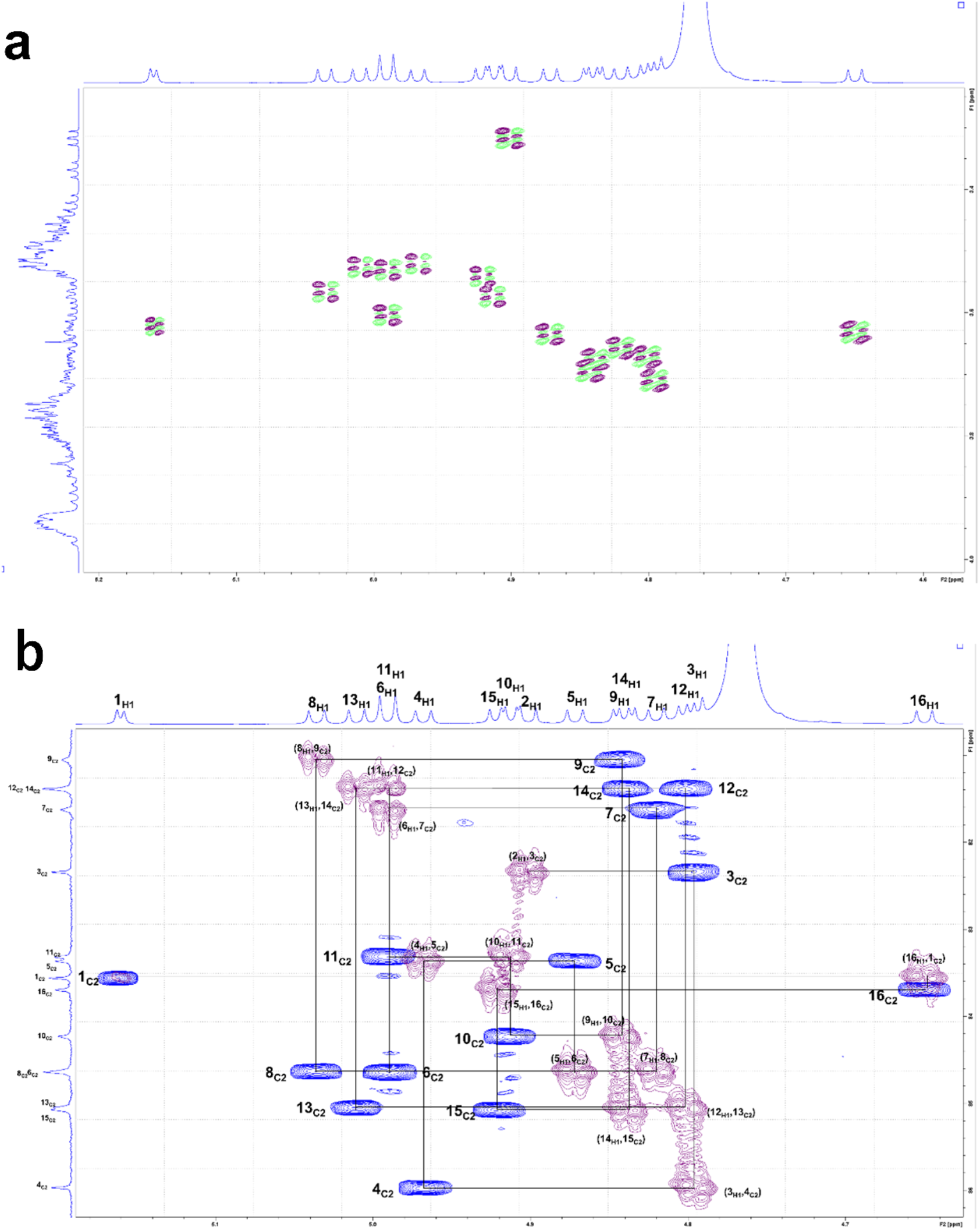
NMR analysis of the β-1,2-glucoside bonds of the main product produced by XccOpgD. **a**, The 2D ^1^H-^1^H COSY spectrum of the main product from XccOpgD. Green and brown peaks represent COSY correlations between H1 and H2. **b**, Overlay of 2D HSQC-TOCSY and HMBC spectra of the main product from XccOpgD. Blue and purple peaks represent HSQC-TOCSY and HMBC correlations, respectively. The number labels beside the peaks are the numbers assigned for Glc moieties in Fig. 1c. Black lines trace from the 1_C2_ (2-carbon at the Glc moiety 1) to 2_H1_ (1-proton at the Glc moiety 2) in the direction to the non-reducing end.

**Extended Data Fig. 3.**
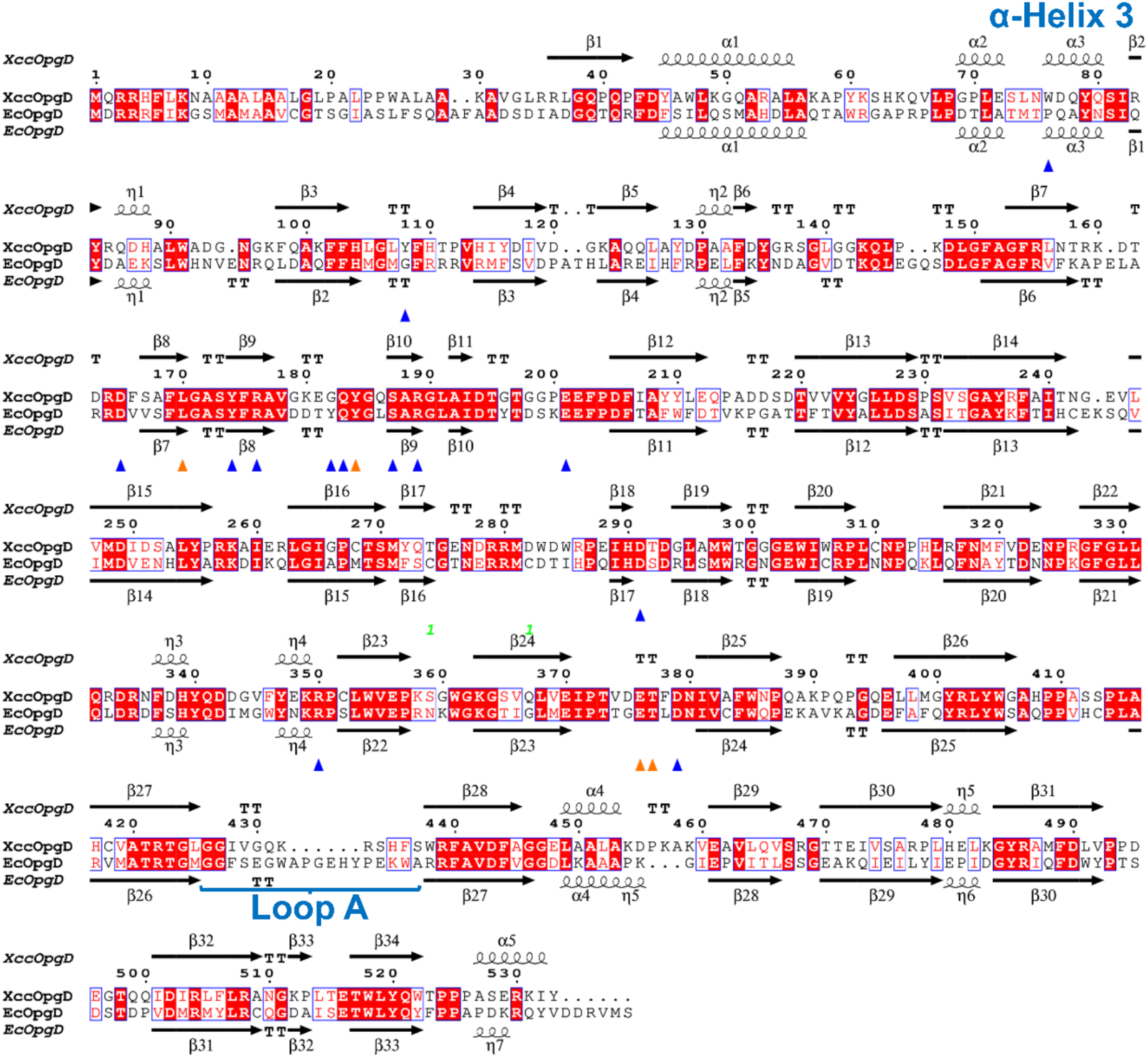
Alignment of XccOpgD and EcOpgD. The alignment was performed using Clustal Omega and visualized using the ESPript 3.0 server (http://espript.ibcp.fr/ESPript/ESPript/). The secondary structures of XccOpgD and EcOpgD are shown above and below the sequences. The regions of α-Helix 3 and Loop A are highlighted in blue letters. Blue and orange triangles represent substrate recognition residues via side chains and only main chains at subsites –7 to +6 of XccOpgD, respectively.

**Extended Data Fig. 4.**
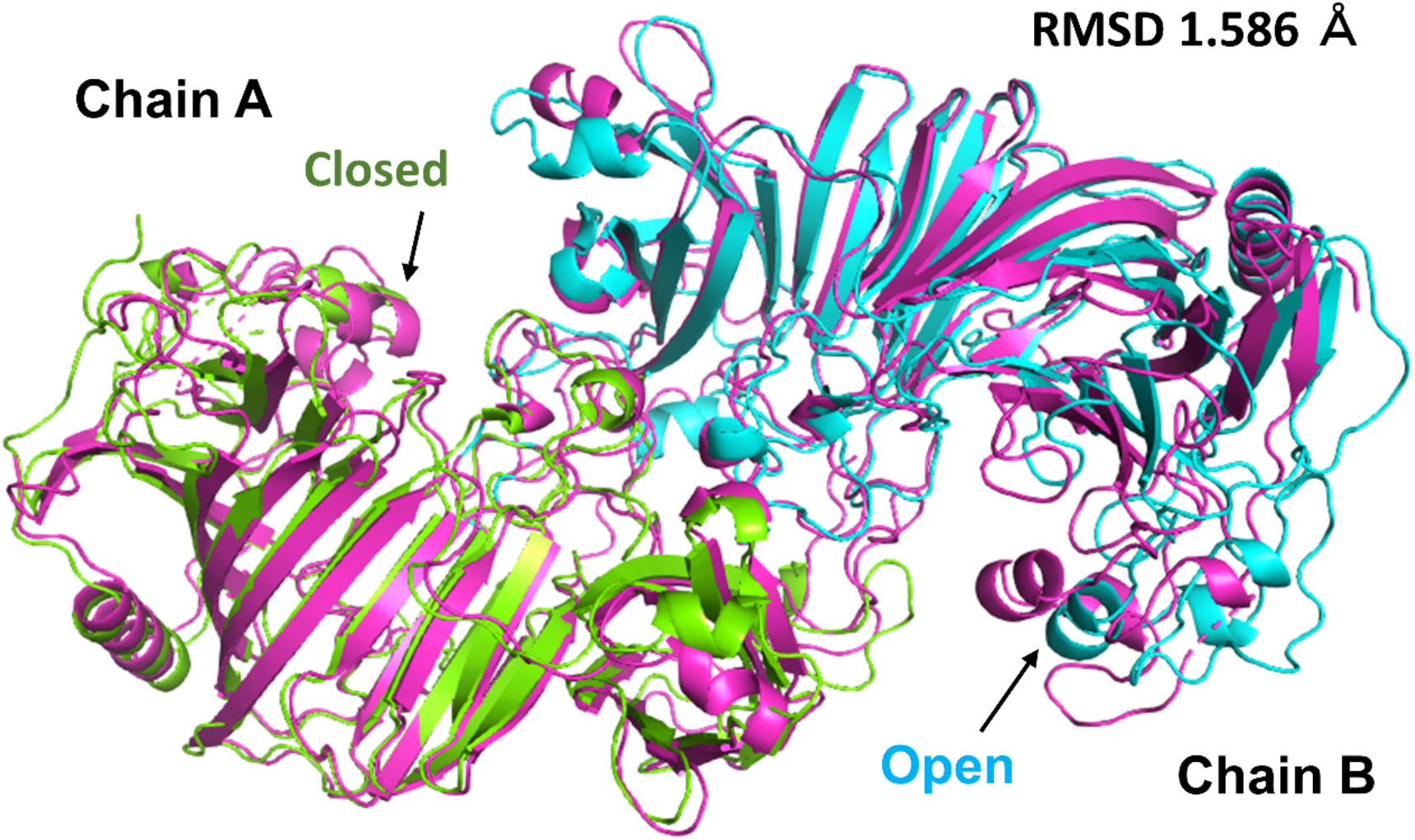
Superposition between Michaelis complexes of XccOpgD and EcOpgD. EcOpgD (PDB ID: 8IP1) is shown as a purple cartoon. Chains A and B of XccOpgD are shown as light green and cyan cartoons, respectively. Substrates are omitted. Chains A and B of EcOpgD form a closed state. In contrast, chains A and B of XccOpgD form closed and open states, respectively, because of crystal packing effects. The root mean square of deviation (RMSD) between XcOpgD and EcOpgD is 1.586 Å.

**Extended Data Fig. 5.**
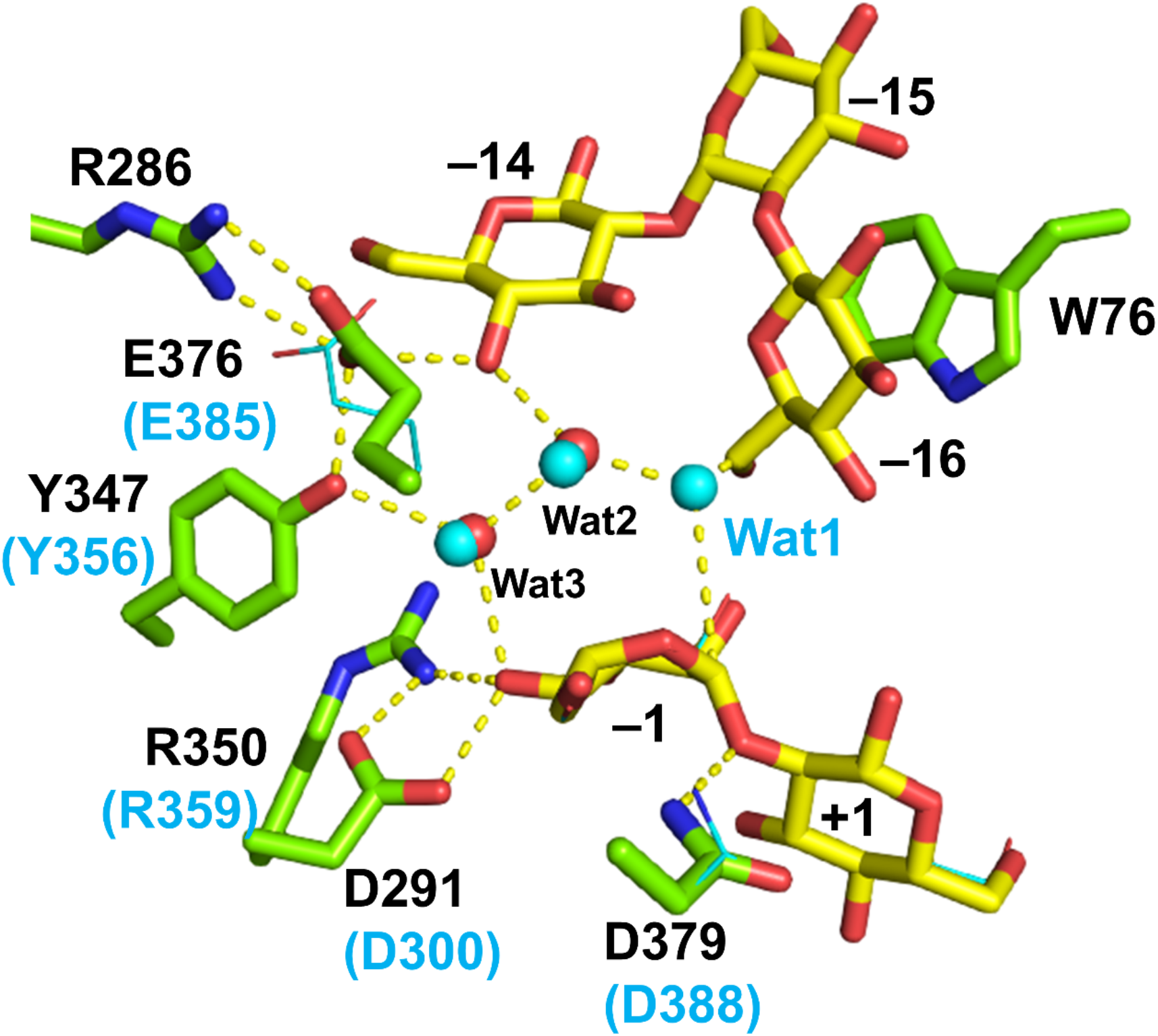
Superimposition between reaction centres of XccOpgD and EcOpgD. Chain A and the substrate of the XccOpgD complex structure (PDB ID: 8X18) are shown as light green and yellow sticks, respectively. Black and cyan labels represent residues in XccOpgD and EcOpgD, respectively. Chain A and water molecules of the superimposed EcOpgD complex structure (PDB ID: 8IP1) are shown as cyan lines and spheres, respectively.

**Extended Data Fig. 6.**
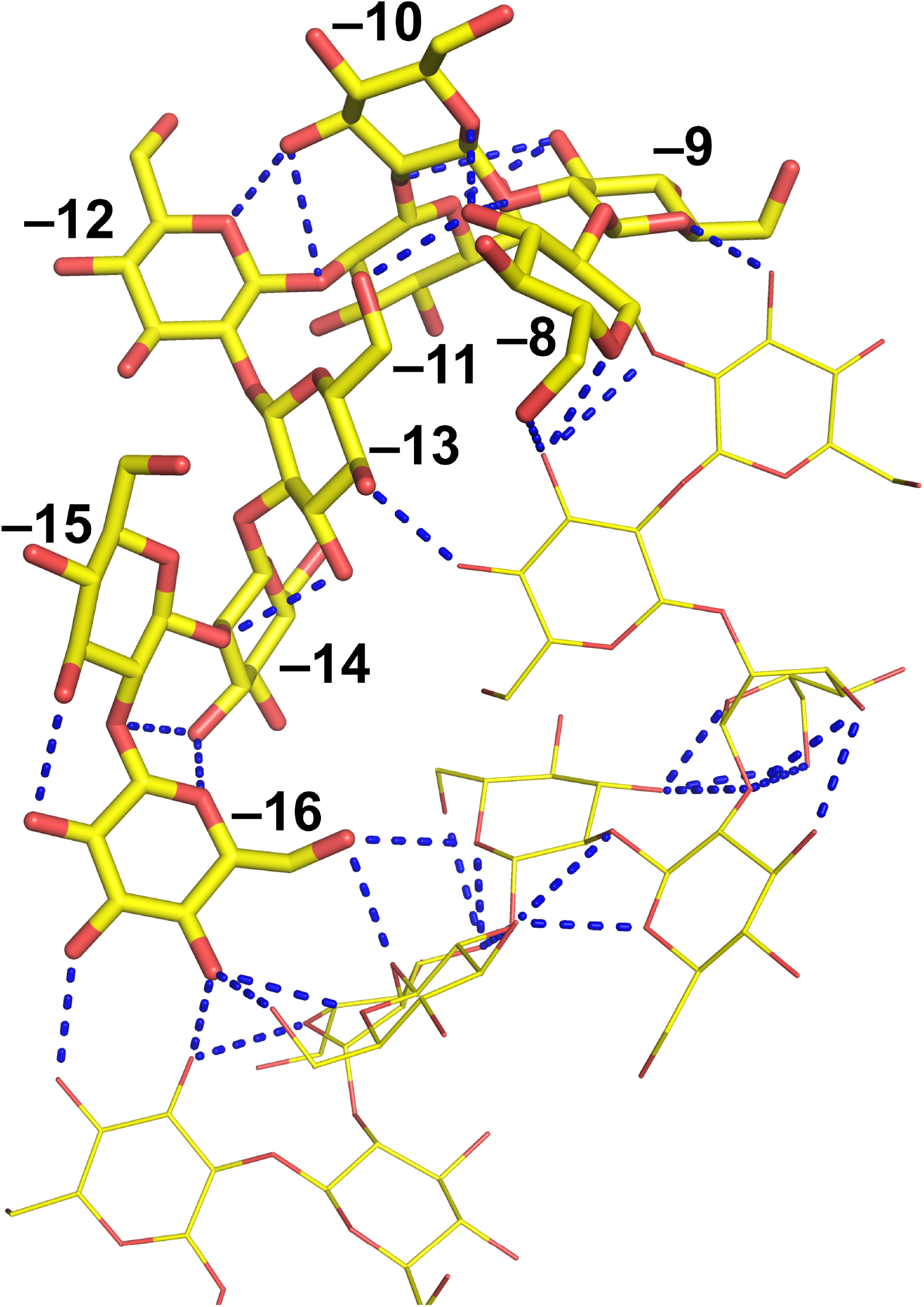
Intramolecular interactions of the β-1,2-glucan at subsites –16 to –8. Substrates of subsites –16 to –8 and –7 to +6 are shown as yellow sticks and lines, respectively. Intramolecular hydrogen bonds are shown as dashed lines. Subsite positions are labelled with numbers.

**Extended Data Fig. 7.**
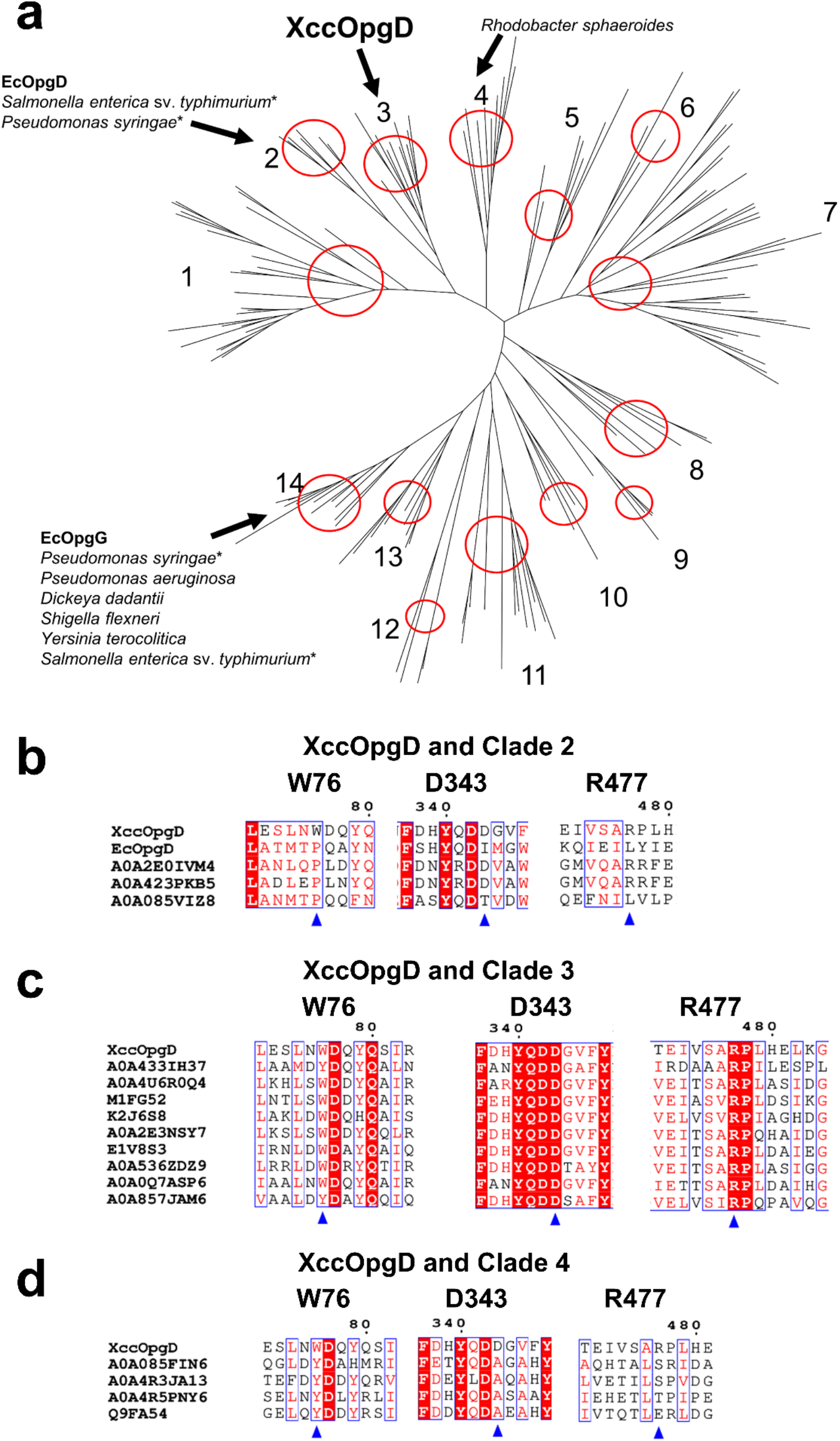
Phylogenetic tree of GH186 and multiple alignments. **a**, Phylogenetic tree of GH186. The figure is taken from Motouchi et al. and partly modified. Each clade is indicated by a red circle with a clade number. Black arrows indicate clades, including homologues from Gram-negative bacteria whose phenotypes of OPG-related gene knockout mutants have been investigated. Asterisks represent species possessing two homologues. **b–d**, Multiple alignments of clades 2–4. The alignments were performed using Clustal Omega (https://www.ebi.ac.uk/Tools/msa/clustalo/) and are visualized using the ESPript 3.0 server (http://espript.ibcp.fr/ESPript/ESPript/). The homologues are represented by UniProt accession numbers. Residue numbers of XccOpgD are shown above the alignment.

