## Supplementary information for "Phytopathogenic cyclic glucohexadecaose from an inverting transglycosylase"

**Supplementary Table 1 | Primer pairs for XccOpgD mutants**

| | Forward primer | Reverse primer |
|---|---|---|
| D379N | accttcaacaacatcgtggcgttctgg | gatgttgttgaaggtctcgtccacggt |
| D291N | atccacaataccgatggcctggcgatg | atcggtattgtggatttctgggcgcca |
| R350A | gagaaggctccgtgcctgtgggtggag | gcacggagccttctcgtagaacacgcc |
| E376Q | gtggaccaaaccttcgacaacatcgtg | gaaggtttggtccacggtggggatctc |
| Y347F | gtgttctttgagaagcgcccgtgcctg | cttctcaaagaacacgccgtcgtcctg |
| T377S | gacgagtctttcgacaacatcgtggcg | gtcgaaagactcgtccacggtggggat |
| W76A | ttgaacgctgatcagtaccagtcgatc | ctgatcagcgttcaaggattccagcgg |
| Q431A | gtgggcgctaaacgcagccatttctcc | gcgtttagcgcccacgatgccgcccag |
| R477A | tcggcggctccgctgcatgagctcaag | cagcggagccgccgagacgatctcggt |
| D342A | taccaggctgacggcgtgttctacgag | gccgtcagcctggtaatgatcgaaatt |
| D343A | caggacgctggcgtgttctacgagaag | cacgccagcgtcctggtaatgatcgaa |
| Cloning (Wild-type) | actttaagaaggagatatacatatggccaaggcagtggggctgcgtcgc | agtggtggtggtggtgctcgaggtagatcttgcgttcgctcgccgg |

All primer pairs are represented from 5′ to 3′.

The forward primer for wild-type was designed to eliminate the N-terminal signal sequence predicted by the signalP6.0 server.

The positions of the mutations are underlined.

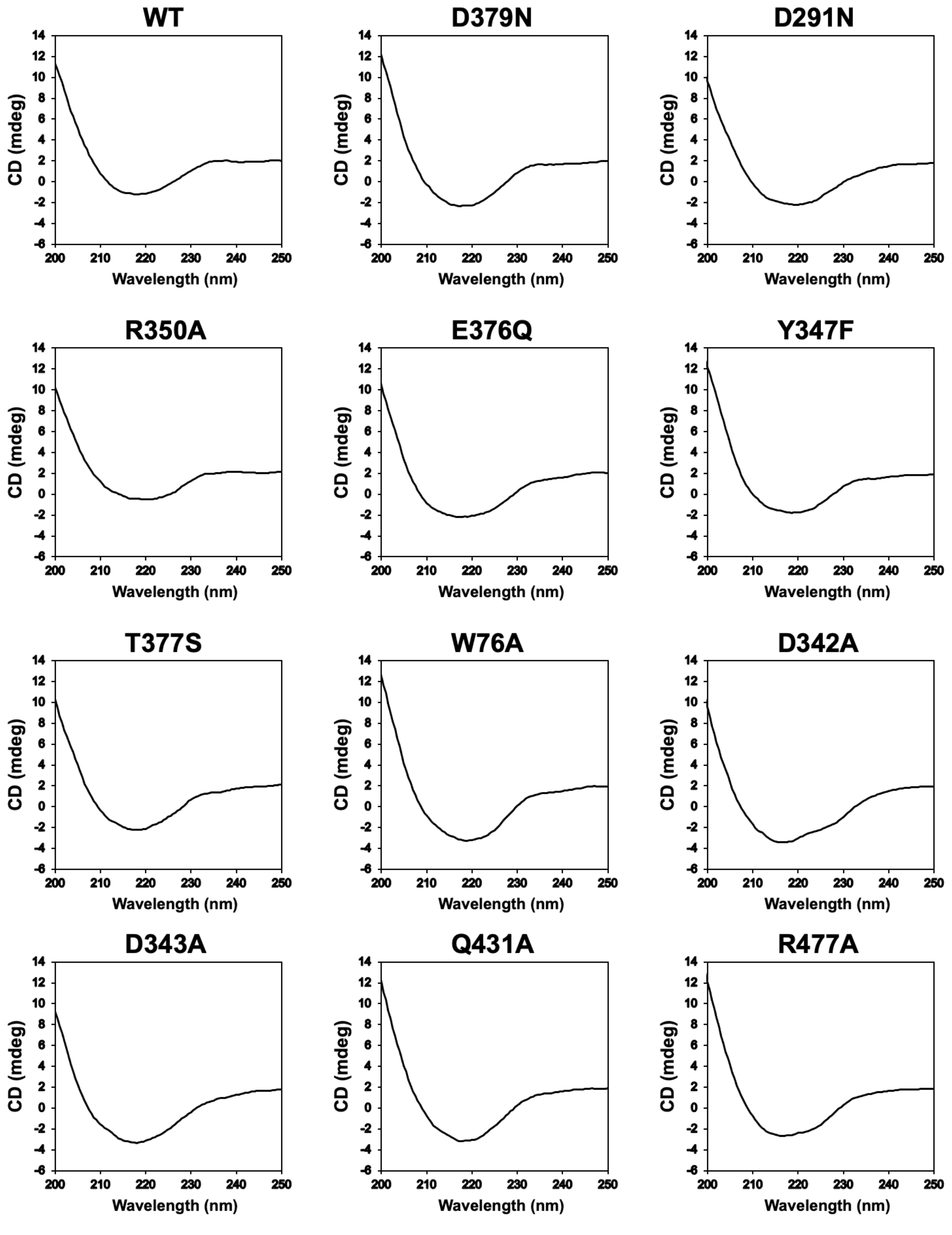


**Supplementary Fig. 1 | CD spectra of wild-type (WT) XccOpgD and mutants.**

**Supplementary Note 1 | The abbreviation for SGL**

The abbreviation BGL for beta-glucanase is indistinguishable from beta-glucosidase. In addition, there appears to be no standard abbreviation for beta-glucanase. Therefore, as an acronym that would be easy to distinguish linkage positions, “S” representing sophoro (-oligosaccharide), an alternative name for β-1,2-gluco-oligosaccharide was adopted as the abbreviation of β-1,2-glucanase.

**Supplementary Note 2 | Molecular mass of products released by XccOpgD**

Molecular masses of cyclic-glucans with DP 32 and 48 (C32 and C48, respectively) were detected by ESI-MS (Fig. 1b). They were also detected when the purified CβG16α was analysed by ESI-MS (Extended Data Fig. 1), indicating that the detection of C32 and C48 is derived from detecting two or three CβG16αs with one or two water molecules. This observation is consistent with the NMR results and Michaelis complex of XccOpgD. In addition, a weak peak likely representing C17 was detected. However, this peak was too weak to confirm the presence of C17 (Fig. 1b). Furthermore, HSQC data demonstrated clearly that the purified product is hexadecaose. Overall, the product of XccOpgD is α-1,6-cyclized β-1,2-hexadecaose.
