## Supplementary figures and images for "Phytopathogenic cyclic glucohexadecaose from an inverting transglycosylase"

### Supplementary Data1

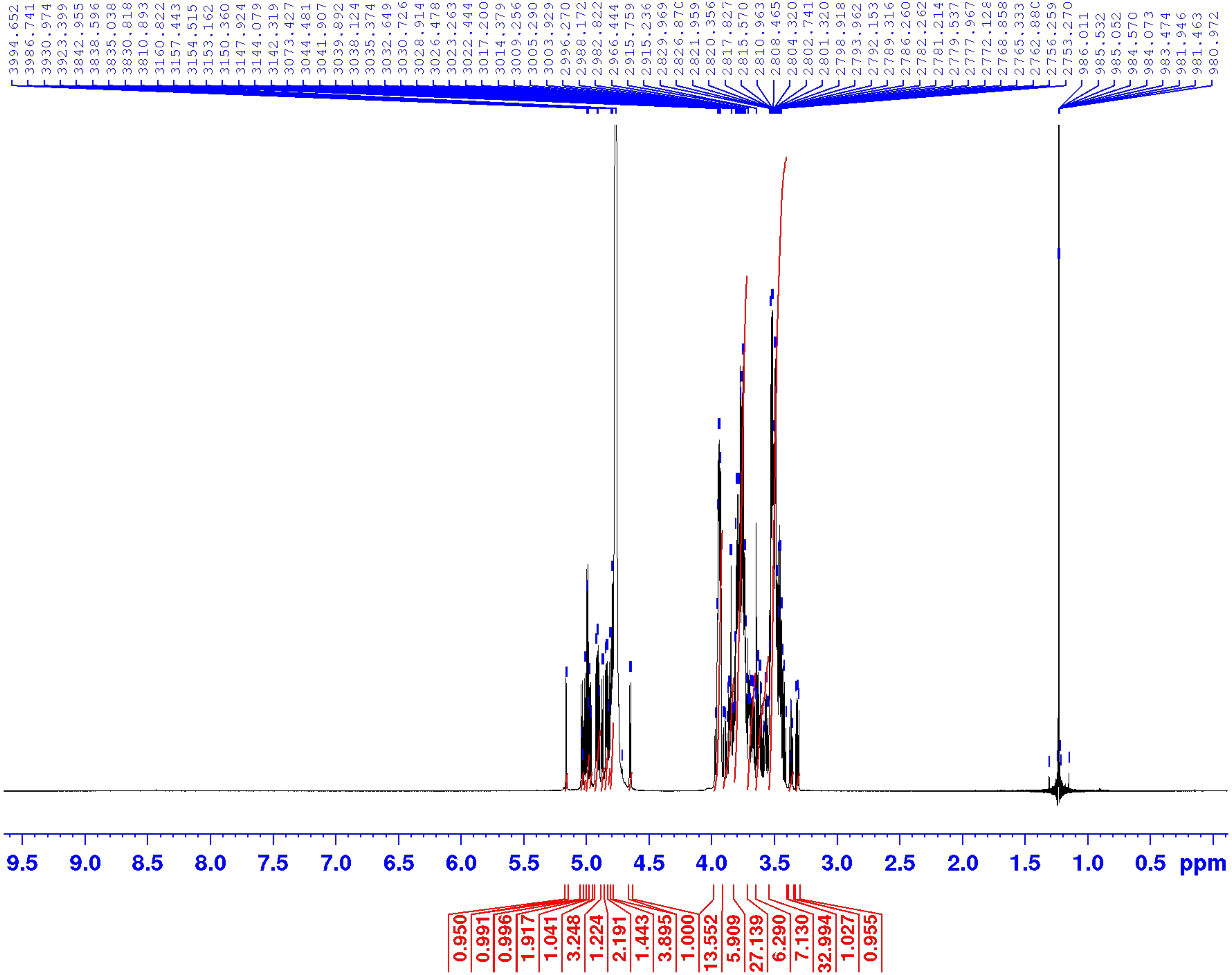

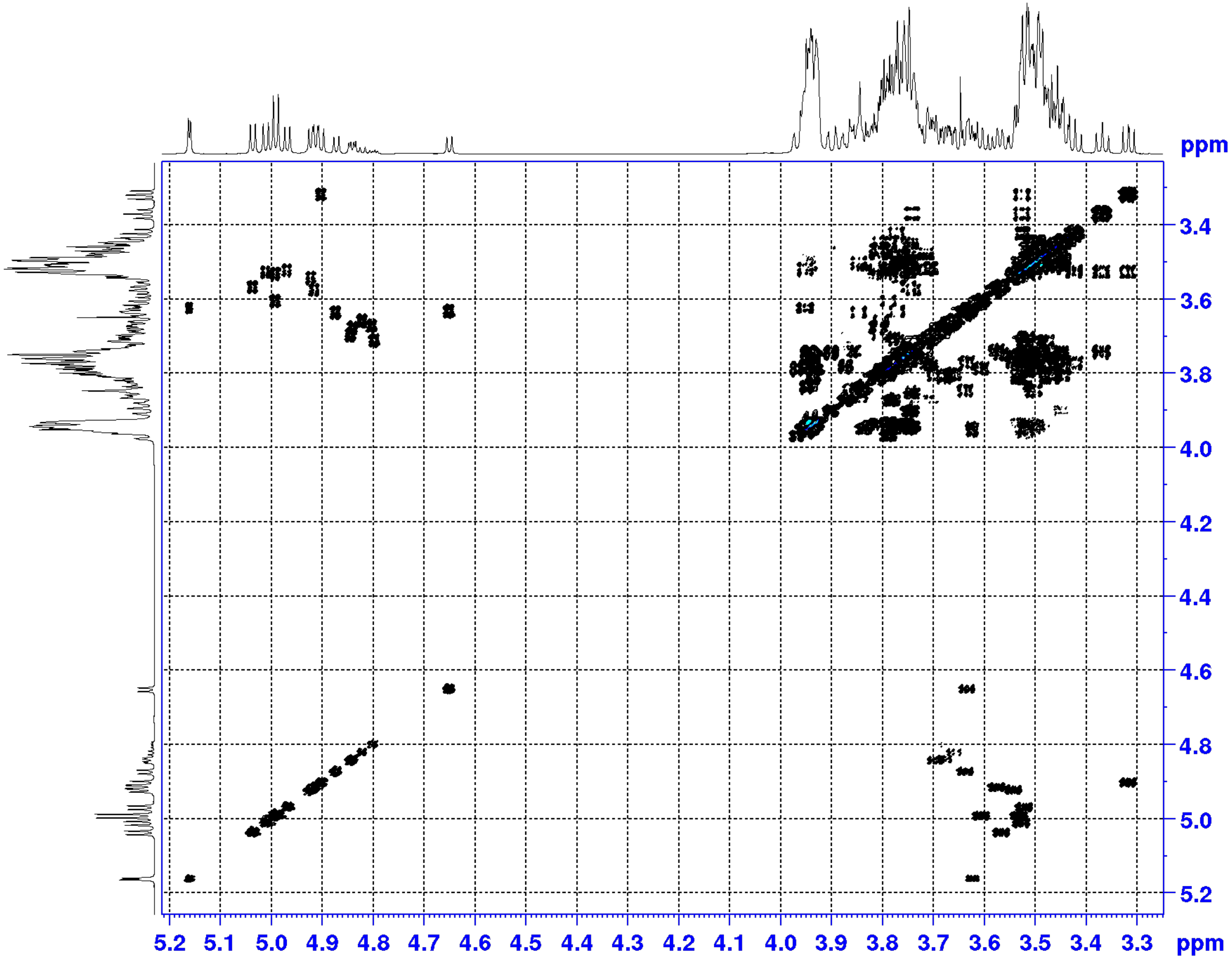

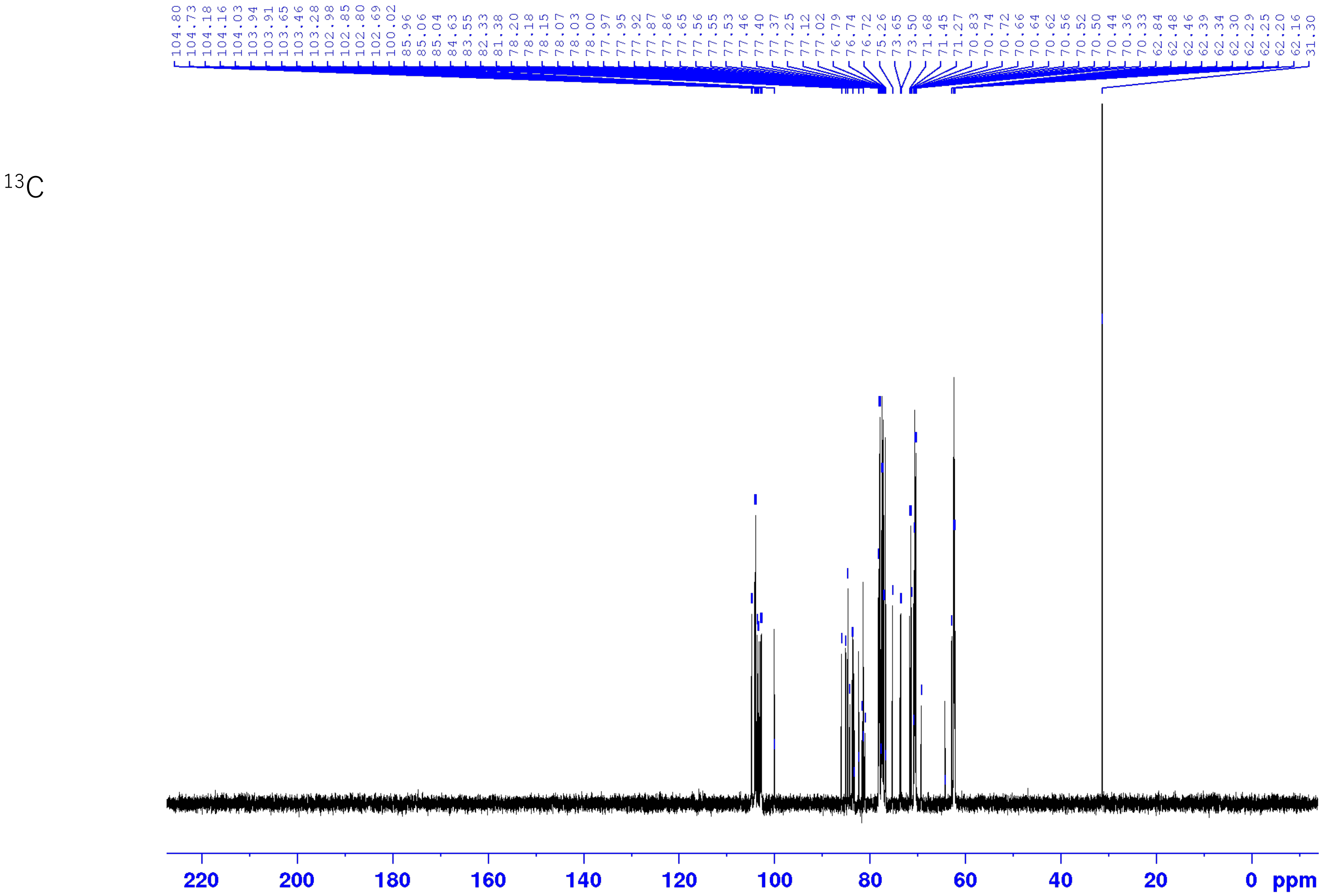

DEPT135

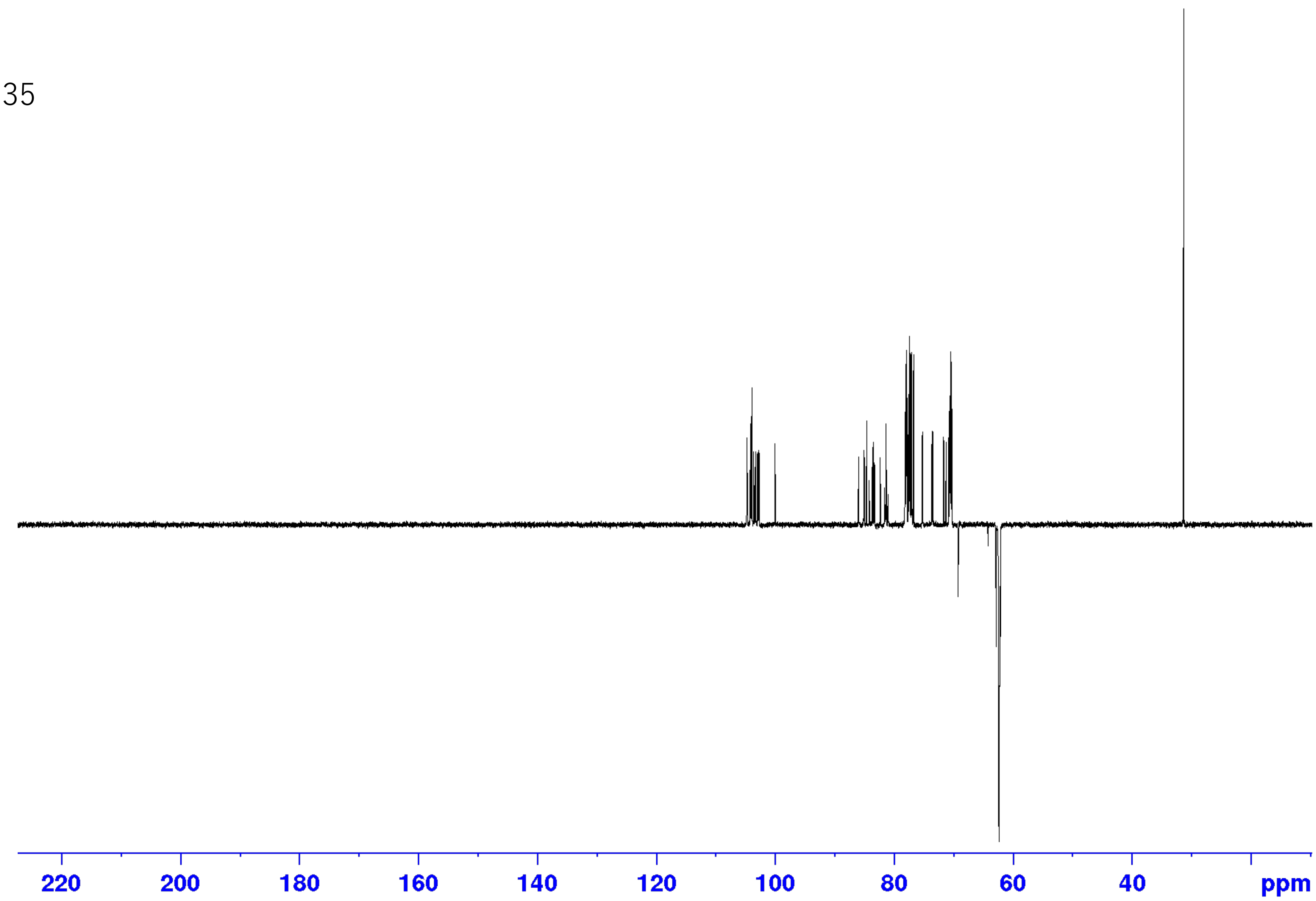

HSQC

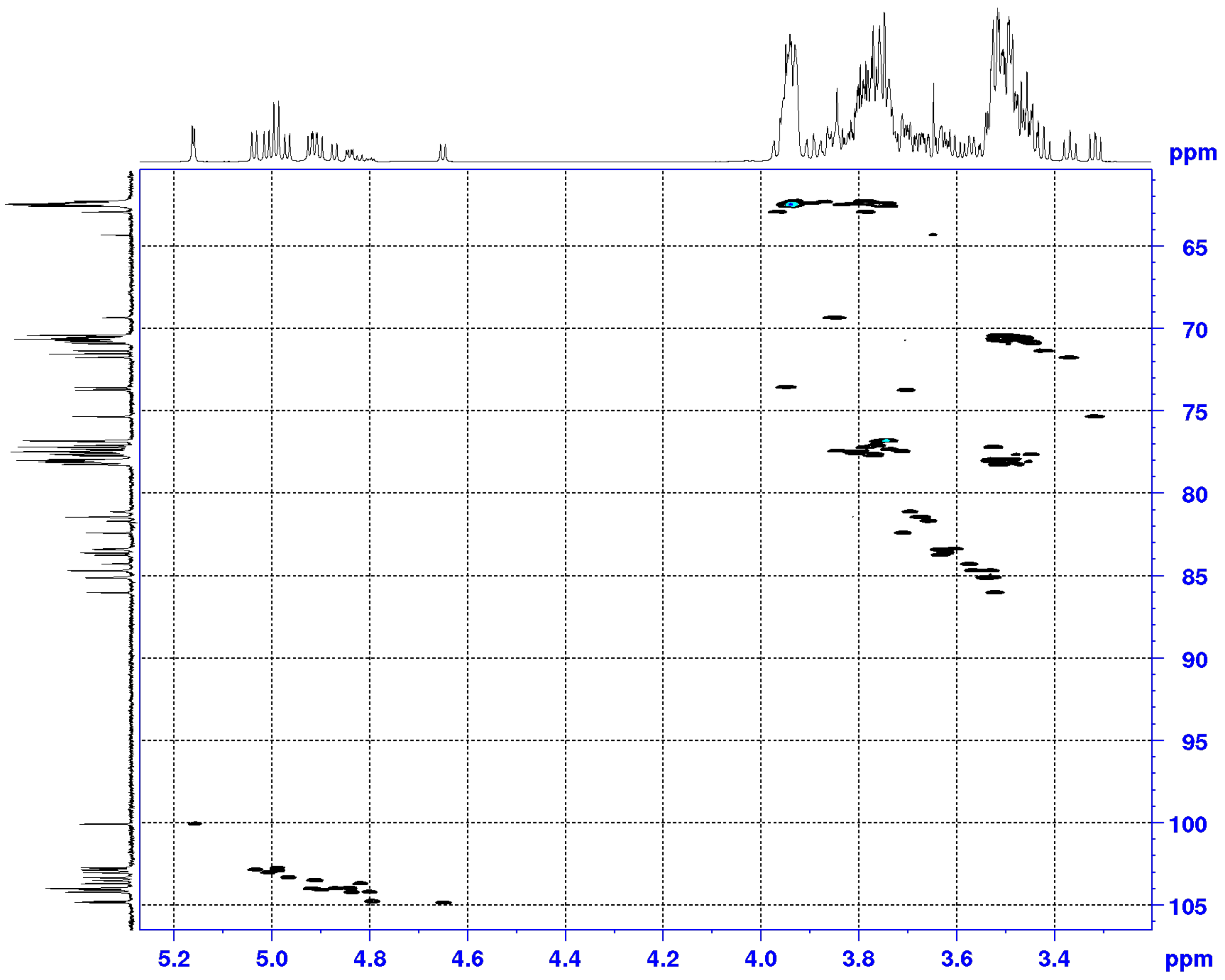

HMBC

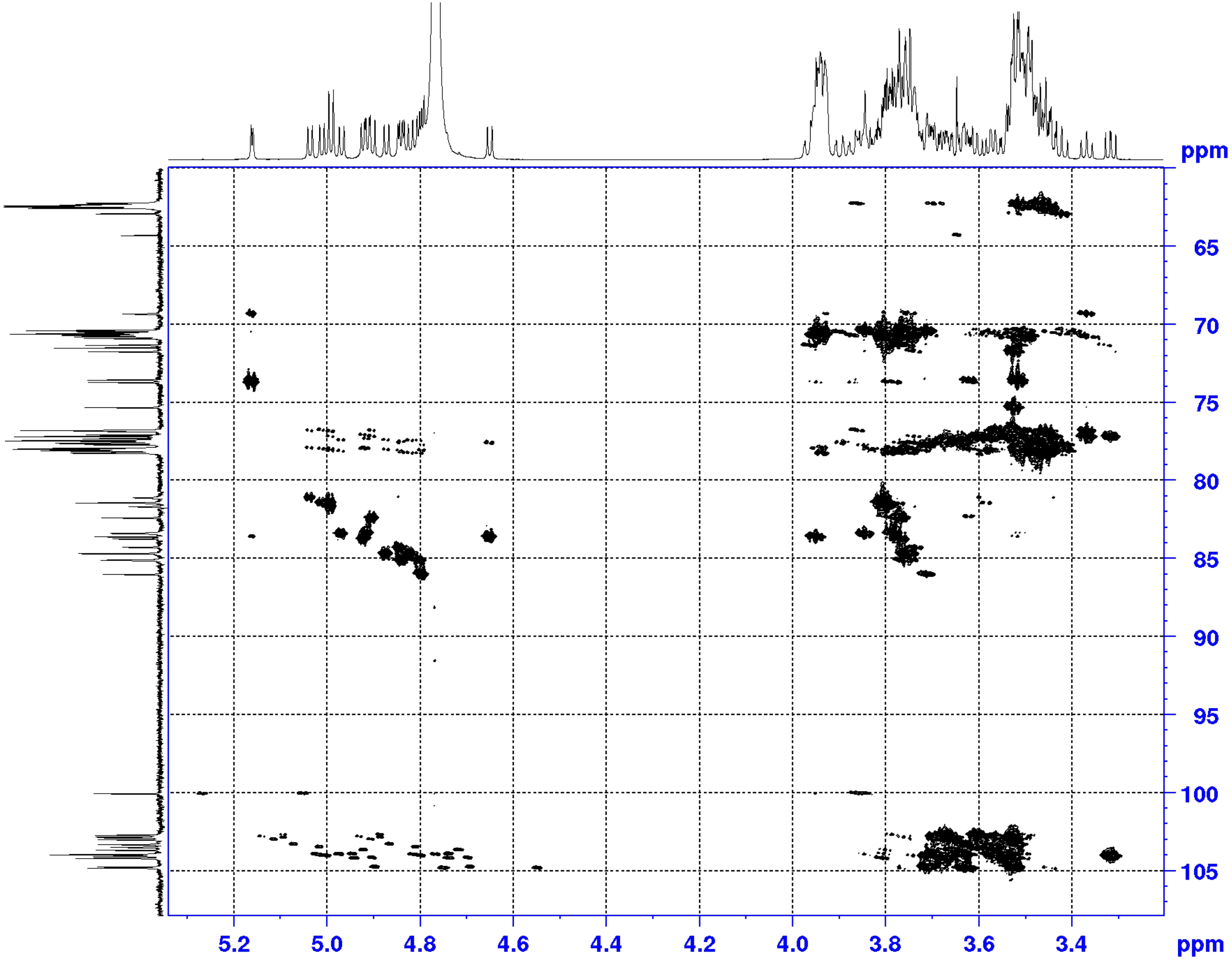

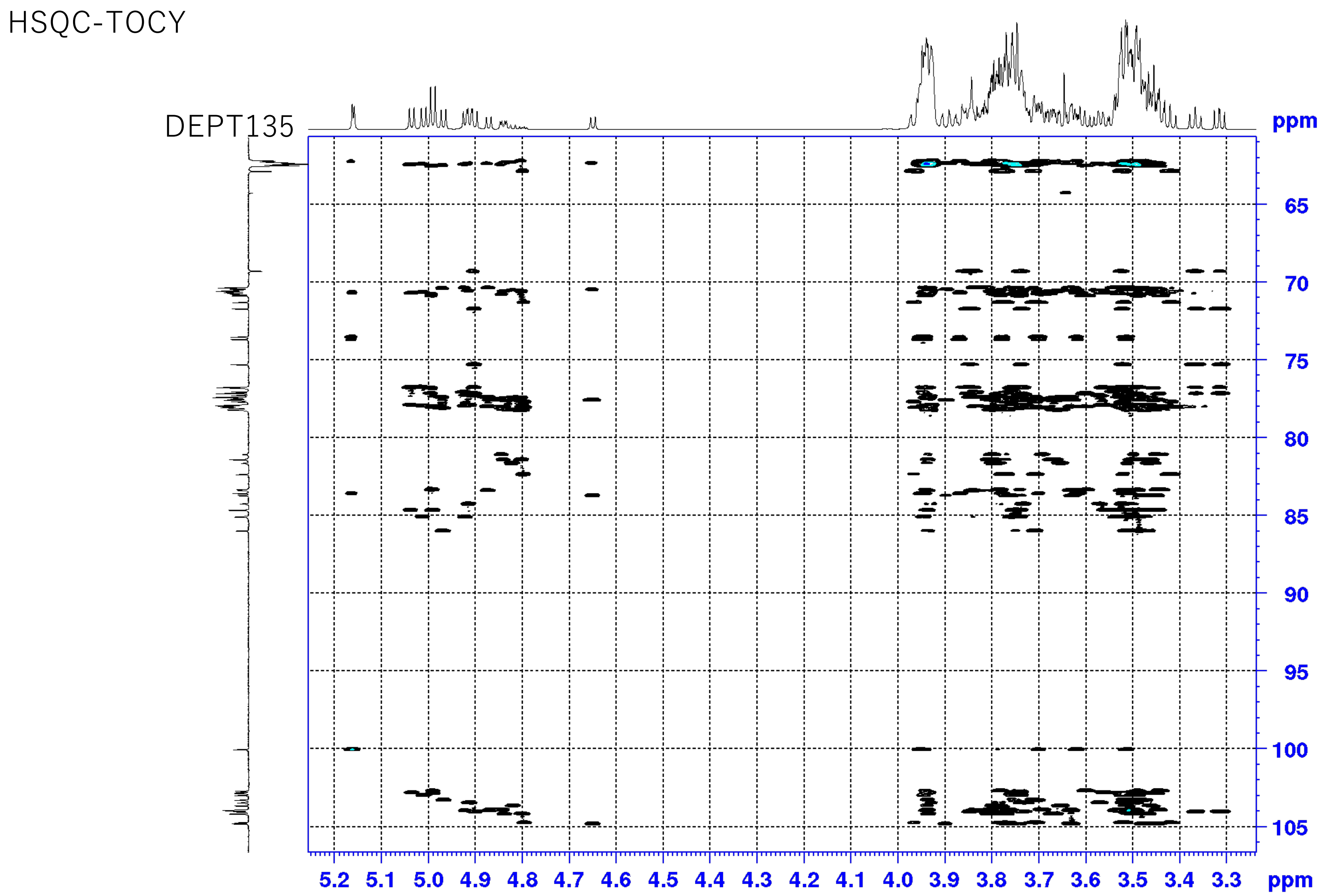
